# A feedback-driven brain organoid platform enables automated maintenance and high-resolution neural activity monitoring

**DOI:** 10.1101/2024.03.15.585237

**Authors:** Kateryna Voitiuk, Spencer T. Seiler, Mirella Pessoa de Melo, Jinghui Geng, Tjitse van der Molen, Sebastian Hernandez, Hunter E. Schweiger, Jess L. Sevetson, David F. Parks, Ash Robbins, Sebastian Torres-Montoya, Drew Ehrlich, Matthew A. T. Elliott, Tal Sharf, David Haussler, Mohammed A. Mostajo-Radji, Sofie R. Salama, Mircea Teodorescu

## Abstract

The analysis of tissue cultures, particularly brain organoids, requires a sophisticated integration and coordination of multiple technologies for monitoring and measuring. We have developed an automated research platform enabling independent devices to achieve collaborative objectives for feedback-driven cell culture studies. Our approach enables continuous, communicative, non-invasive interactions within an Internet of Things (IoT) architecture among various sensing and actuation devices, achieving precisely timed control of *in vitro* biological experiments. The framework integrates microfluidics, electrophysiology, and imaging devices to maintain cerebral cortex organoids while measuring their neuronal activity. The organoids are cultured in custom, 3D-printed chambers affixed to commercial microelectrode arrays. Periodic feeding is achieved using programmable microfluidic pumps. We developed a computer vision fluid volume estimator used as feedback to rectify deviations in microfluidic perfusion during media feeding/aspiration cycles. We validated the system with a set of 7-day studies of mouse cerebral cortex organoids, comparing manual and automated protocols. The automated protocols were validated in maintaining robust neural activity throughout the experiment. The automated system enabled hourly electrophysiology recordings for the 7-day studies. Median neural unit firing rates increased for every sample and dynamic patterns of organoid firing rates were revealed by high-frequency recordings. Surprisingly, feeding did not affect firing rate. Furthermore, performing media exchange during a recording showed no acute effects on firing rate, enabling the use of this automated platform for reagent screening studies.

## Introduction

Recently, advances in biological research have been greatly influenced by the development of organoids, a specialized form of 3D cell culture. Created from pluripotent stem cells, organoids are effective *in vitro* models in replicating the structure and progression of organ development, providing an exceptional tool for studying the complexities of biology [1]. Among these, cerebral cortex organoids (hereafter “organoid”) have become particularly instrumental in providing valuable insights into brain formation [2–4], function [5, 6], and pathology [7, 8]. Despite their potential, organoid experiments present significant challenges. Brain organoids require a rigorous, months-long developmental process, demanding substantial resources and meticulous care to yield valuable data on aspects of biology such as neural unit electrophysiology [9], cytoarchitecture [10], and transcriptional regulation [8].

The primary methods for generating and measuring organoids depend on media manipulations, imaging, and electrophysiological measurements [11], which are all labor- and skill-intensive, limiting the power and throughput of experiments [12]. Cell culture feeding and data collection occur at intervals realistic for researchers. Furthermore, during manual feeding and data collection, the cell cultures are removed from the incubator, which provides a controlled gas, temperature, and humidity environment [13]. Ideally, feeding should be aligned with the cells’ metabolic cycles, and data should be collected at intervals on par with the biological phenomenon. The disturbance incurred by leaving the incubator environment is shown to increase metabolic stress and batch-to-batch variability, potentially impacting the quality of the experiment [14], as well as increasing contamination risk. These limitations hinder the depth of insights gained from these organoid models, particularly in studies focused on dynamic neural processes and disease modeling [11].

Laboratory robotics, most often liquid handling devices [15], offer increased precision and throughput but are primarily designed for pharmaceutical screens, limiting their adoption in research labs due to high costs, large footprints, and inflexible workflows [16]. Moreover, many of these systems lack the ability to seamlessly integrate new technologies as they emerge. Conversely, academic research labs are benefiting from advancements in commercial and custom-made technologies, facilitated by in-house fabrication methods like 3D printing [17, 18], which are enhancing their capacity to manipulate and measure biological systems. However, without an easyto-integrate, device-agnostic robotic platform, researchers are constrained to manual operations, restricting the power and scope of their experiments. By outfitting devices to carry out automated jobs and relay data through communication networks, they acquire around-the-clock functionality and increased fidelity [19]. The flexibility in size (number of devices per integrated system) allows researchers to optimize for the experimental design and budget. Implementing programmable feedback loops derives precision and self-optimization by dynamically adjusting to real-time data [20–22], offering a practical alternative to complex mathematical modeling for experiment control. This approach would enable more integrated, flexible automation in research settings, broadening the scope and efficiency of experiments.

Automating multiple devices to report data presents a challenge for device management and communication, necessitating flexible and efficient infrastructure. Addressing this need for an interconnected ecosystem of devices, services, and technologies is possible through designing networks using standards defined by the Internet of Things (IoT). This approach has already impacted wearables [23], agriculture [24], city infrastructure [25], security [26], and healthcare [27]. It was recently proposed to expand this approach to biology research [28]. Previously, each researcher built a custom device and code from scratch with unique assumptions for communication and behavior. Each device operated in solitude, lacking integration and feedback with other devices. Here, we establish a platform that addresses these challenges, combining electrophysiology, microscopy, microfluidics, and feedback control, automated and integrated through IoT technology for touch-free, in-incubator tissue research.

## Results

### An integrated microfluidic, electrophysiology, and imaging organoid research platform

We have developed an integrated platform (Figure 1) that automates organoid culture and data collection in individual microenvironments. While microfluidics (Figure 1a) controls the media environment, digital microscopy captures the morphogenic features. The neural activity is recorded by local field potential measurements using complementary-metal-oxide semiconductor (CMOS) high-density microelectrode arrays (HD-MEA)[29](Figure 1b). The IoT cloud network brokers the communication between all devices and facilitates data storage, processing, and presentation services including an interactive webpage (Figure 1d). Through touch-free automation, samples remain undisrupted in the incubator, increasing the consistency of images and allowing for higher frequencies of feeding and recording.

**Fig. 1.**
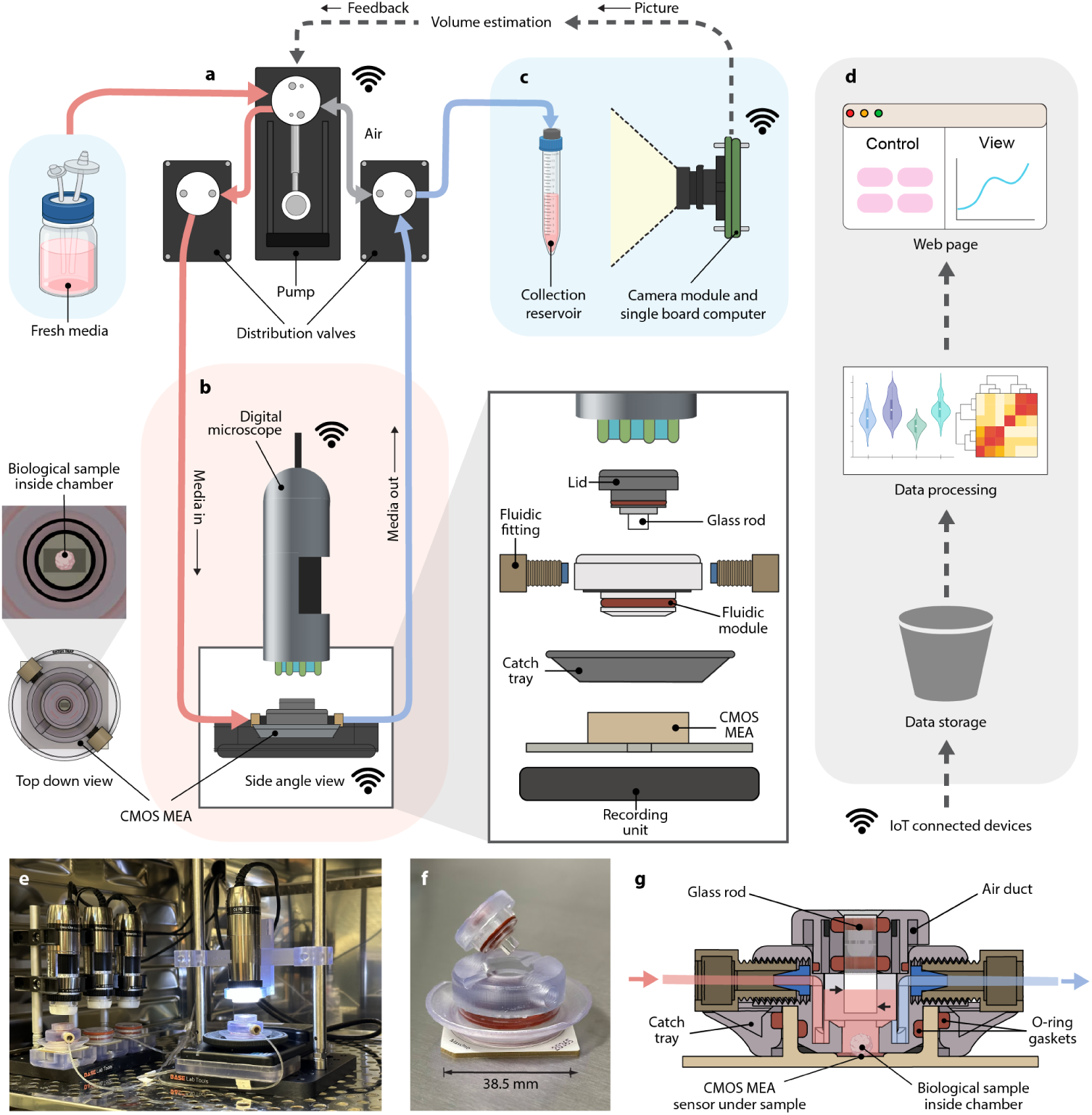
Schematic diagram of the integrated feedback platform. **(a)** A syringe pump and valve system dispense fresh media and aspirate conditioned media at user-defined intervals. The blue background represents 4°C refrigeration. **(b)** Microscopy and HD-MEA electrophysiology record morphology and functional dynamics of the biological sample. The red background represents 37°C incubation. Exploded view: the microfluidic culture chamber for media exchange couples with the HD-MEA. **(c)** A camera captures images of the aspirated conditioned media drawn from each culture and relays them through cloud-based data processing for volume estimation feedback to the syringe pump system. **(d)** Devices communicate over MQTT (Message Queuing Telemetry Transport) protocol and automatically upload data to the cloud, where it is stored, processed, and presented on a web page. **(e)** The experimental setup in the incubator shows two microfluidic culture chambers and two conventional membrane lids. **(f-g)** 3D printed microfluidic culture chamber and cross-section diagram. The media level, noted by the upper black arrow (559µL) and lower black arrow (354µL) on the glass rod, is the ideal operating range that keeps the rod immersed in media. The biological sample is adhered to the HD-MEA sensor at the bottom.

At user-defined intervals, conditioned media is aspirated by a syringe pump through a system of distribution valves (Figure 1a), stored in a collection reservoir (without passing through the syringe pump vial) (Figure 1c), and replaced by an equivalent volume of fresh media. Both types of media are perfused through flexible fluorinated ethylene propylene (FEP) tubing at 110 mm/s, which leads to low shear forces [30] (see Materials and Methods, Microfluidic cell culture). This equates to a flow rate of 44.1 µL/s.

The digital microscope (Figure 1e) is attached using 3D-printed parts on aluminum posts. The 3D printed culture chambers integrate the microfluidics and HD-MEAs. A liquid-impermeable O-ring gasket ensures media retention inside the chamber. The well lid includes a polished glass rod submerged in the media, improving image quality and removing the effects of condensation. Alignment grooves in the glass rod lid prevent rotation and incorrect fitting. The lid exchanges gas with the incubator conditions through ventilating air ducts (Figure 1g), similar to a cell culture well plate. The removable and re-attachable lid reduces manufacturing complexity and enables future use of other lids with applications beyond imaging.

Figure 1c shows the cross-section of the culture chamber attached to the HD-MEA. The media flows in (red) and out (blue). The sinuous media path and well geometry ensure minimum disturbance to the biological sample [30]. Fresh media is delivered on top of the volume present in the chamber, similar to partial media changes found in manual feeding protocols [31, 32]. The ideal operating range is between 350 to 700 µL (see Supplementary Materials and Methods, Figure 1 and Table 1 for numerical volume limits). In the case of over-aspiration, media drops to a minimum of 170 µL before aspirating air from the chamber’s headspace. The 3D-printed catch tray guards against overflow, collecting up to 1.5ml (200% of the chamber’s capacity) to protect the recording equipment from liquid damage.

### Computer vision for microfluidic flow feedback

We developed a computer vision volume estimation system to monitor the accumulation of aspirated media and identify anomalies during culture feeding events. Figure 2a-b provides a detailed view of the setup inside a refrigerator, which includes three main components: a collection reservoir support system, an LED panel, and a camera module (see Materials and Methods, Assembled devices and custom 3D-printed components). The camera module remains on standby for image capture requests made by other IoT devices or users. Upon request, computer vision techniques are employed to estimate the media volume within the reservoirs accurately.

**Fig. 2.**
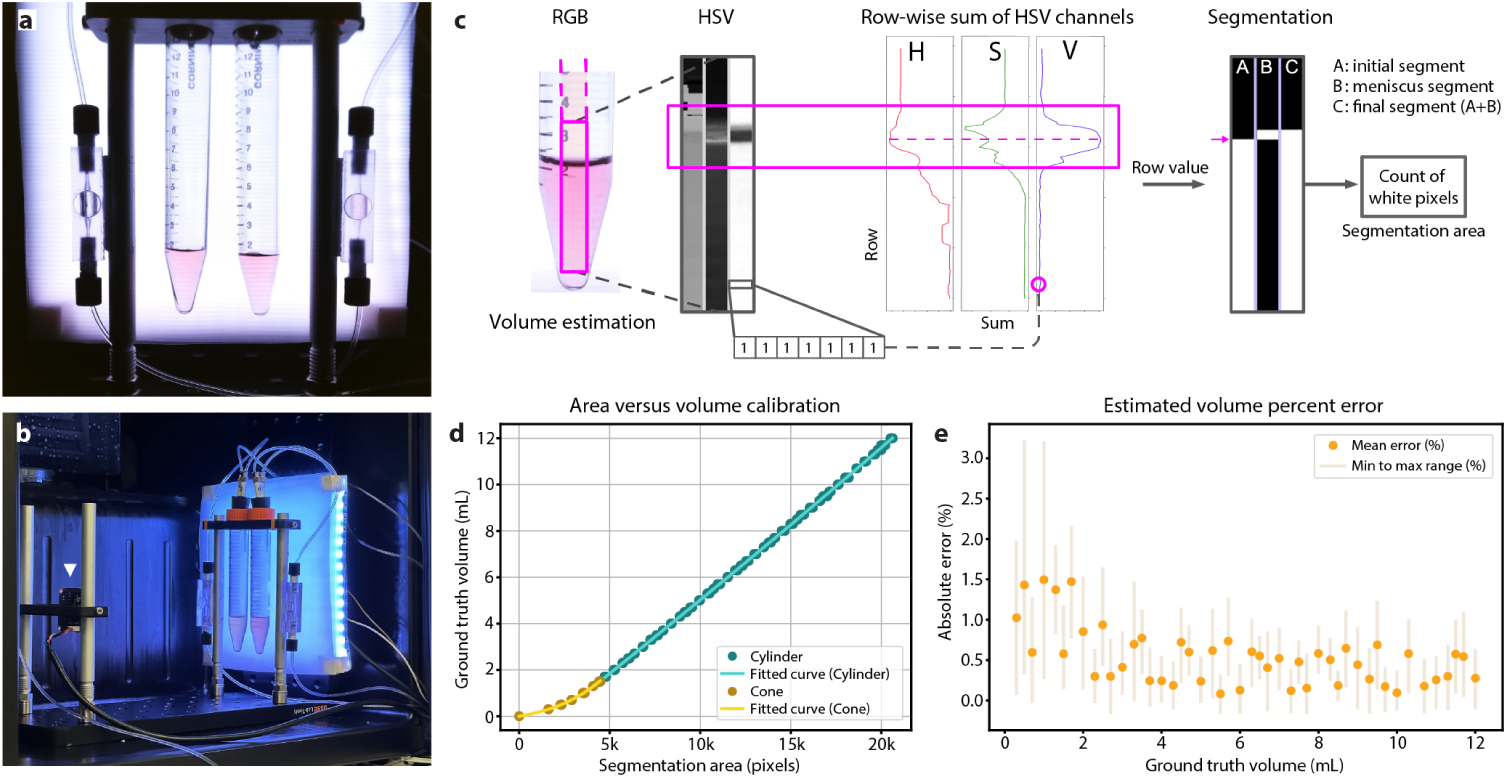
Computer vision for volume estimation. **(a)** Example of a raw image captured by the camera module. **(b)** In-refrigerator volume estimation setup in Figure 1c. The CMOS camera module (the white triangle) images the conical tubes with a diffused LED backlight for even illumination. **(c)** Fluid segmentation: a rectangular pixel patch down the center of the conical tube; Row-wise summations of the HSV channels are used to detect the location of the meniscus. The initial liquid potion segmentation is added to the meniscus portion to yield the final segmentation. **(d)** Calibration graph with a fitted relationship of segmented pixel count to ground truth volume. **(e)** The absolute error percentage: orange dots represent the average error at selected volumes. The shaded bar represents the minimum to maximum error range.

Figure 2c shows the computer vision process (see Materials and Methods, Computer vision for fluid volume estimation) for segmenting area related to the media in the reservoir. A calibration was required to establish the relationship between the segmented area in pixels and volume in milliliters. We captured 184 images of the collection reservoirs containing volumes of media ranging from 0 to 12 mL (several pictures for each volume), with each volume confirmed by a scale, accurate to 1 µL. For each specific volume in Figure 2d, multiple points overlap and are all accounted for to calculate the polynomial regression lines. To accommodate the reservoir’s conical section (volumes *<*1.5 mL) and cylindrical section (volumes *>*1.5 mL), two distinct regressions were applied, ensuring a high degree of precision for each geometrical shape.

A Leave-One-Out cross-validation (LOO) [33] approach was employed to quantify the model’s error. This method tests the model’s accuracy and generalizability in an unbiased manner, ensuring that the calibration results in a model that performs reliably across different samples. The effectiveness of the model is assessed quantitatively with the following metrics: an average Mean Absolute Error (MAE) of 0.56% (equivalent to 27 µL), an average standard deviation of errors at 0.53% (22 µL), and an average Root Mean Square Error (RMSE) of 0.77% (35 µL). The polynomial models exhibit R-squared values of *>*0.99, denoting an optimal fit of pixel area to liquid volume. Figure 2e shows the average absolute error percentage at a specific volume, with the bar indicating the error range from minimum to maximum.

### IoT infrastructure creates an ecosystem of devices and cloud-based services

We built a cloud-based IoT ecosystem that enables communication between users, devices, and services to implement actions, record data, and streamline upload, storage, and analysis. All devices (here: pumps, microscopes, and microelectrode arrays) run software using the *device-class* Python framework (Figure 3a and Supplementary Materials and Methods). Devices operate collectively with shared core software and complementary behaviors: they can request jobs from each other, yield during sensitive operations, and ensure collaborative functions and smooth operation (Figure 3d). Devices update their *shadow* in the database whenever their state information changes (i.e, assigned experiment, schedule, current job and estimated completion time, and other dynamic variables) to eliminate the need for device polling. Messages (i.e., job requests) between devices and services are sent through a centralized MQTT broker via the publish/subscribe protocol. This decoupled architecture allows for independent and extensible deployment of components. Data generated by devices is immediately uploaded to an S3 object storage in a predefined structure using an experiment Universally Unique IDentifier (UUID) as the top-level key. A ‘metadata.json’ file stores experiment details, sample information, notes, and an index of the produced data. Raw data is stored separately from analyzed data under different sub-keys. Cloud jobs, which operate as shared services, process raw data from S3 and write results back to S3, reporting status via MQTT messages. To utilize the IoT ecosystem, users initiate experiments, control devices, and visualize data through a website (see Materials and Methods, Website and screenshots in Supplemental Figure 2), with the typical user workflow in Figure 3c.

**Fig. 3.**
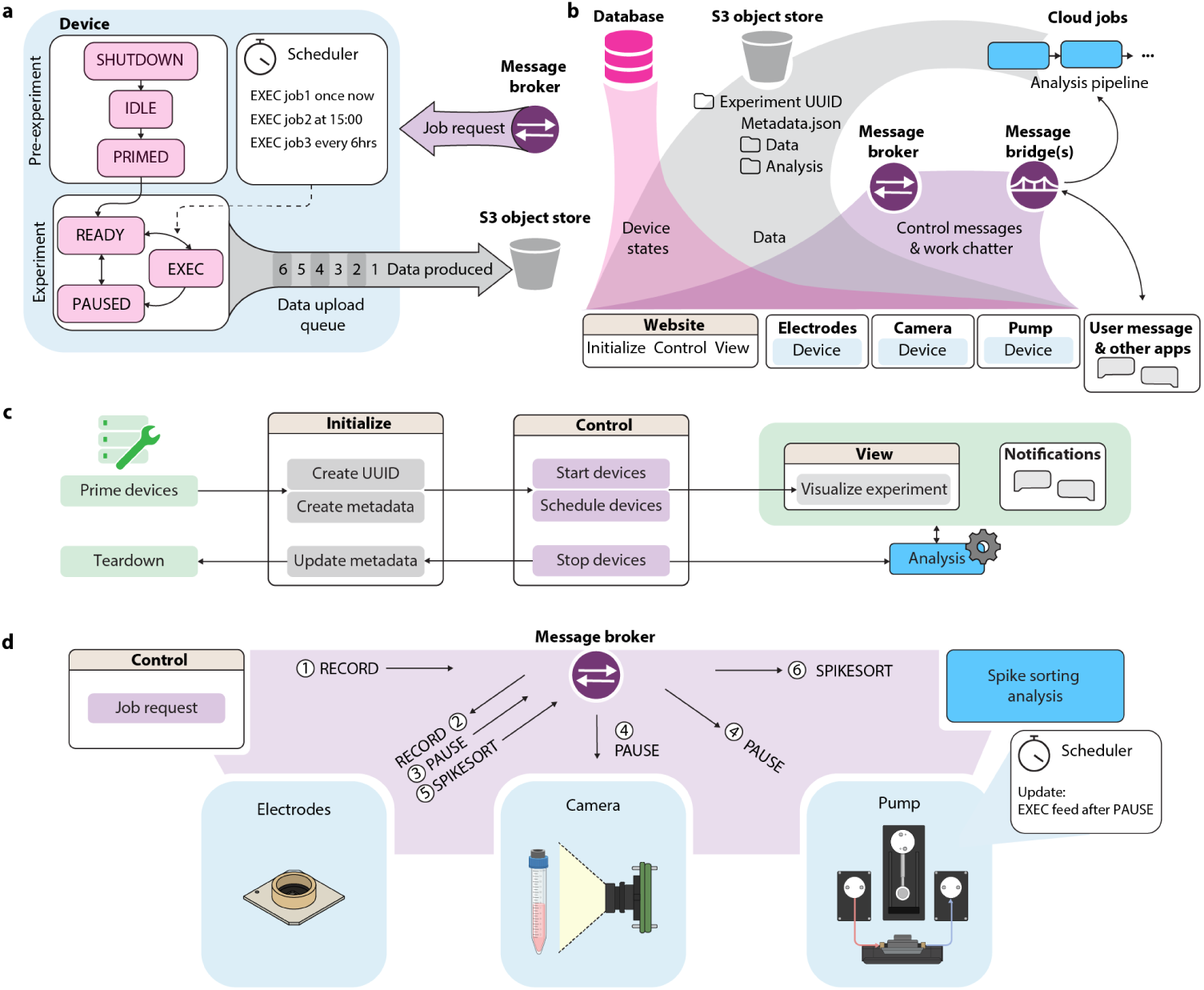
Cloud-based device interactions. **(a)** The *device-class* is a generalized state machine framework of all IoT devices. The *device* participates in experiments by taking in job requests (from experimenters or other devices), scheduling and executing the jobs, and producing data files that are queued and uploaded to cloud storage. **(b)** IoT infrastructure. Device states (pink) are saved in a database and displayed on the website user interface. Device-generated data (gray) is saved and organized in cloud storage, where it can be accessed by user interface or analysis cloud jobs. Devices send communications (purple) through a message broker and use message bridges to translate messages to analysis pipelines or text messaging applications. **(c)** User workflow. Devices are physically primed in accordance with experimental procedures such as sterilization. On the ‘Initialize’ webpage, an experiment is created with a unique ID (UUID) and descriptive notes (metadata). On the ‘Control’ webpage, devices are called to start working on the experiment and are given a job schedule. The ‘View’ webpage and notifications allow the user to monitor the ongoing experiment. **(d)** Example of inter-device communication: (1) A RECORD job request is made from the ‘Control’ panel. (2) The message broker delivers the record request to the electrophysiology recording unit. (3) The electrophysiology unit pauses all other devices to ensure a quality recording. (4) All devices receive a pause request. The pump reschedules a feed until after the pause. (5) Upon finishing the recording, the electrophysiology unit delivers a spike sorting request to commence data analysis.

### Automated study of cerebral cortex organoids

The integrated research platform was used to study the effects of automation on the neuronal activity of pluripotent stem-cell-derived mouse cerebral cortex organoids. Embryonic stem cells were aggregated, patterned, and expanded to generate organoids using a previously defined differentiation protocol [34, 35]. Day 32 post-aggregation, organoids were plated two-per-chip directly onto HD-MEAs. Two were plated to maximize use of the HD-MEA surface (3.85 x 2.10 *mm*^2^). For the 7-day study, 8 chips across two batches were split into groups that were fed and recorded with standard manual procedures (Controls, N=4), manual feeding and automated recording (AR, N=1), automatic feeding and manual recording (AF, N=1), or automatic feeding and automatic recording (AFAR, N=2). All chips were imaged in the incubator every hour, each using a dedicated upright digital microscope (DinoLite).

Automated microfluidic feeds were used to increase the consistency and frequency of cell culture media replacement. We removed conditioned cell supernatant from the well and dispensed the equivalent volume of fresh media for each feed cycle. The controls had 1.0 mL media replacement every 48 hours, consistent with standard protocols. AF and AFAR were placed on a protocol in which 143 µL media were replaced every 6 hours, matching the total media volume turnover across groups for the 7-day study. The schedule of automated media feeds was defined at the experiment’s launch and initiated by a timed feeding job command sent to the microfluidic pump. The fidelity of feeding was controlled through a computer vision volumetric feedback loop on the aspirated conditioned media (Figure 2, 4a).

Conditioned media has a high protein content, contains cellular debris, and is susceptible to forming salt crystals [36, 37]. In microfluidic systems, this leads to clogs, error accumulation, and failure modes [38]. To overcome this, a volume estimation feedback loop was initiated each time the pump performed a job. At the time the medium was perfused to/from a specific well, the pump sent a job request to the camera module responsible for imaging the well’s collection reservoir. The image was captured, uploaded to the cloud, its volume estimated by the computer vision Estimator, and returned to the pump for feedback interpretation. Within tolerance, the action was declared a success (marked as a green check mark in Figure 4a), and no further action was taken. Outside of tolerance, the pump scheduled itself a new job proportional to the volume discrepancy and in relation to the number of previous feedback attempts (see Materials and Methods, Feedback interpreter).

**Fig. 4.**
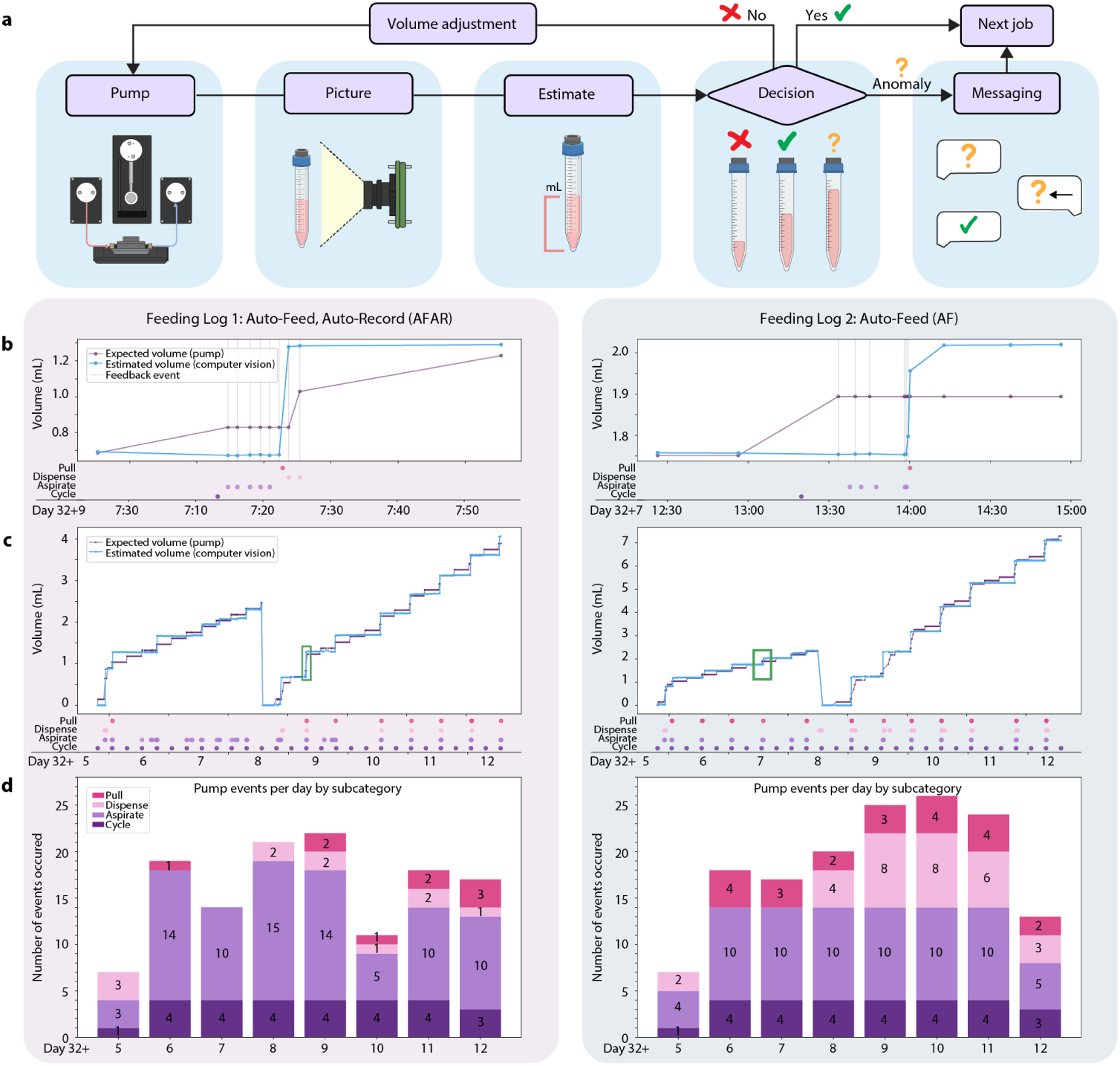
Volume feedback. **(a)** Volume estimation feedback loop. After the pump completes a microfluidic action, it requests a picture of the media collection reservoir from the camera module. The picture is passed to the cloud-based computer vision program to estimate the current volume. The results is compared with the expected volume, and a decision is made: within tolerance (green checkmark), a microfluidic volume adjustment action is needed (red “x”), or an anomaly is detected (yellow question mark). Once the estimated volume is within tolerance (green check mark), the feedback cycle ends and proceeds to the next job. If this cannot be achieved or an anomaly is detected, such as out-of-range volumes, an alert is sent to the user messaging service to request assistance. **(b-d)** On these graphs, the “Day” x-axis summarizes the timeline: organoids were plated on the HD-MEA on Day 32, automation started 5 days after plating and continued to day 12. Above this axis, dots mark the occurrence of microfluidic events. **(b)** Graphs of the Expected Volume and Estimated Volume for the automated AFAR 1 (left) and AF (right) during a period of feedback events. Event types are marked with dots below the graph. **(c)** The complete view of Expected and Estimated volume traces over the 7-day study. **(d)** Stacked histogram pump events per day organized by type.

The system was designed to resolve discrepancies using feedback. However, in cases where volume estimation returns a value outside of reason (i.e., *>* expectation + 2 mL) or if the feedback iteration limit is reached (i.e., *>* 20 attempts), the system was programmed to send an alert to a Slack messaging channel and pause. During both batches of the 7-day experiments, the system resolved errors independently, and this condition was not reached.

The automated feeding and feedback results for AF and AFAR 1 are visually represented in Figure 4b-d. Figure 4c shows the traces of expected volume and computer vision estimated volume for AFAR 1 (left) and AF (right) for the 7-day study (Days 5 to 12 post-plating). There was a collection reservoir change on Day 8 in which the 15 mL conical was replaced with a fresh tube. In both samples, the drop in estimated and expectation reflects the collection reservoir exchange. For AFAR 1 (Figure 4b, left), a zoomed-in view of the feedback loop following the scheduled feeding cycle at 7:12 on Day 9 highlights feedback actions taken to remedy a volumetric discrepancy. In this instance, the volume estimation was less than expected after the feed cycle. Five consecutive aspiration jobs were carried out, and the estimated volume still remained under expectation. At the 6th iteration of feedback, a pull job was sent to the pumps, which raised the collection volume above the expected volume. In the 7th and 8th iterations of feedback, two dispense jobs were engaged to supplement the well for the over-aspiration. In a similar case, for AF (Figure 4b, right), a total of 6 iterations of feedback were engaged to bring the estimated volume into tolerance with the expected volume; however, in this example, no dispense jobs were required. Figure 4d shows histograms of the sum of pump events per day by subcategory. Each feeding cycle (four per day) was scheduled, and all other events occurred through feedback.

### High-frequency HD-MEA recordings and automated feeding do not disrupt neuronal activity

To evaluate organoid neuronal activity, extracellular field potentials were measured using 26,400 electrode HD-MEAs, which can record up to 1,020 electrodes simulta-maps derived from the first and final activity scans for each sample are presented in Figure 5b, with an outline of the organoid edge based on alignment of corresponding microscopy image. To optimize electrode coverage, we generated specific configuration files for electrode selection based on the regions with the highest activity, which remained constant for seven of the eight chips. In one case (AF, Day 32+6), we updated the configuration due to the emergence of a new high-signal area on the second day of recording. Stable configuration maps allowed for automated electrode recordings over days, optimizing long-term analysis.

**Fig. 5.**
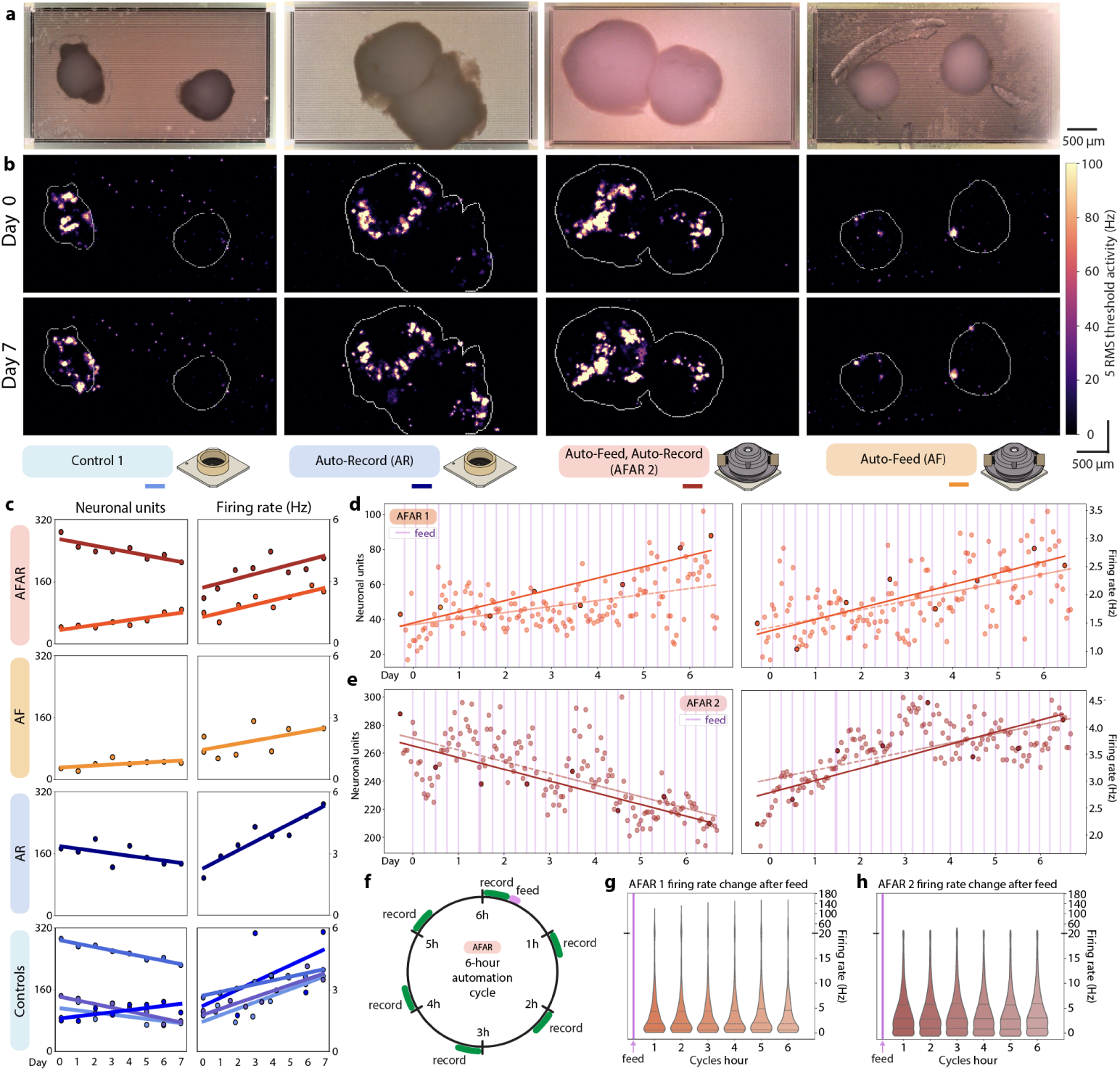
Electrophysiology analysis of the 7-day cerebral cortex organoid study. **(a)** Digital microscope images of example organoid conditions. **(b)** Boundaries of each organoid were outlined using image segmentation and overlaid with activity scans from the initial recording “Day 0” (top) and last recording “Day 7” (bottom). Experimental conditions are labeled underneath, with a color legend. **(c)** Total detected neural units (left column) and median firing rates per unit (right column) in daily 10-minute recordings, grouped by experimental condition. **(d-e)** Detected neural units (left) and median firing rates (right) in hourly resolution for AFAR 1 **(d)** and AFAR 2 **(e)** using their automated 10-minute recordings. Feeding events are noted as vertical violet lines. **(f)** The two AFAR samples had a 6-hour automation cycle that included one 143 µL feed (violet) and six 10-minute recordings (green). **(g-h)** Violin graph of all unit’s firing rates per recording for AFAR 1 **(g)** and AFAR 2 **(h)** organized into bins of the 6-hour automation cycle following each feeding event. The 6-hour feeding regimen did not induce cyclical changes for either sample.

Manual recordings involved an experimenter placing each HD-MEA on the recording unit and initiating 10-minute recordings via software. In contrast, the hourly recordings (AFAR 1-2 and AR) featured the HD-MEA remaining on the recording unit while automated software handled the entire process, from power management to data uploading. AFAR 1-2 and AR each amassed over 158 recordings, totaling over 26.3 hours (525 GB) of electrophysiology data per sample. Conversely, all manually recorded samples (Controls 1-4 and AF) accumulated 7 recordings each, amounting to 1.2 hours (23 GB) of electrophysiology data. The studies generated over 2 TB of data from over 100 hours of mouse organoid recordings.

From these data, we analyzed the effects of our automated microfluidic, imaging, and recording system on the neuronal activity of the brain organoids housed therein. Imaging of the chips from above (Figure 5a) allowed us to align the body of the organoid with neural activity (Figure 5b). In some instances, such as in Control 1, neurite outgrowths were evident in the images and activity scans.

Initial activity scans were used to distribute samples into experimental and control conditions. We sought even distribution of the samples into conditions to reduce biases with respect to initial activity levels. Neural units identified from initial recordings by spike sorting with Kilosort2 [39] were: Control 1: 87 units, Control 2: 292 units, Control 3: 144 units, Control 4: 80 units, AR: 173 units, AFAR 1: 43, AFAR 2: 250, AF: 29. Three chips were omitted from the study for having a unit count less than 25. Throughout the 7-day study, all samples with unit counts 29 to 80 (Control 4, AFAR 1, and AF) increased their detected unit counts during the 7-day study, while all samples with unit counts 87 and above (Control 1-3, AR, and AFAR 2) decreased their detected unit counts (Figure 5c).

The average firing rate per neural unit increased for every sample, irrespective of feeding or recording schedules (Figure 5c). All samples started with an average firing rate of 1.95 Hz (*σ* = 0.48), increased by 0.24 Hz per day (*σ* = 0.09), and concluded with an average of 3.98 Hz (*σ* = 1.22). The automated conditions (AR, AFAR, and AF) presented no divergence from the controls on neural unit count, firing rate, or morphology resulting from increased frequency of feeding and/or increased frequency of recording.

### High-frequency HD-MEA recordings reveal dynamic neuronal activity states in organoids

The hourly recorded conditions (AFAR 1-2 and AR) revealed transient states, not apparent with single daily recordings (Figure 5c-e). Linear regression trendlines comparing the hourly and daily recordings for a single sample are congruent, however daily recordings do not capture the prominent oscillatory dynamics of neuronal unit count and median firing rate captured by the hourly recordings. Median firing rates were observed to fluctuate as much as 3-fold over the course of a day and are not well-characterized by linear regression fit (AFAR 1 *R*^2^=0.31, AFAR 2 *R*^2^=0.42, AR *R*^2^=0.69).

We inspected the effect of feeding on these dynamics. The AFAR samples had a six-hour automation cycle (Figure 5f) that included one 143 µL feed and six 10-minute recordings. We examined effect by aligning recordings to a six-hour ‘time since feed’ cycle. Figures 5g and h present the composite graphs of aggregated neuronal firing rates of AFAR 1 and AFAR 2, comprising 26 feeding cycles with all recordings binned with respect to their time since feeding. The superimposed feeding cycles did not show a trend in units (not shown) nor firing rate in relation to feeding cycles. The variance presented in Figures 5d and e do not align with the six-hour feeding cycle.

### Effects of feeding during recording

We further investigated the effect of feeding with a follow-up experiment that included microfluidic feeds during electrophysiology recordings to capture the immediate response on neuronal activity to a media injection. The AR sample from the 7-day study (Figure 5) was equipped with a microfluidic culture chamber and set on a new feeding and recording schedule. Automated 15-minute recordings were performed every hour for 36 hours. Feedings occurred every third hour that began at minute 5 of the ongoing recording (Figure 6a). Each feeding cycle was defined as an aspiration and dispense of 150 µL. Automated feeds increased in their number of cycles each third hour for four experimental conditions (1 cycle = 150 µL, 2 cycles = 300 µL, 4 cycles = 600 µL, and 6 cycles = 900 µL). Each of these conditions were performed three times (Figure 6b).

**Fig. 6.**
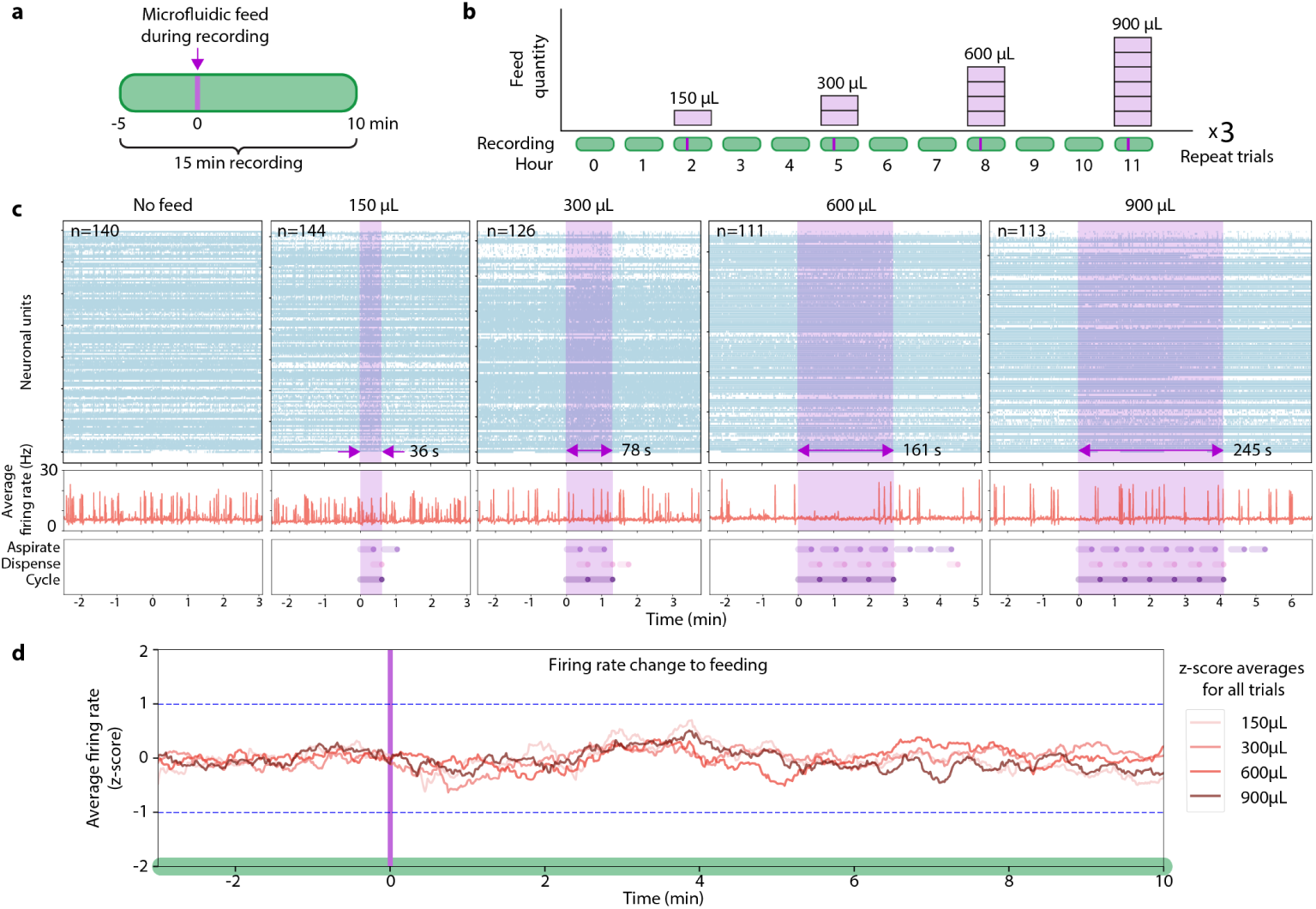
Effect of feeding during recording. **(a)** 15-minute recordings were collected hourly from a cerebral cortex organoid, with automated microfluidic feeding beginning at minute 5. **(b)** Feedings occurred every third hour of ascending fixed-volume cycles of 150 µL, 300 µL, 600 µL, and 900 µL, then repeated for three trials (36 hours). **(c)** Graphs of select recordings showcasing each of the five conditions. Top: spike raster of neuronal units (y axis) firing over time (x axis). Middle: average firing rate per neuronal unit computed by dividing the total spikes in a 100 ms bin by number of neuronal units. Bottom: microfluidic feeding actions performed by the pump. Each feeding “cycle” is composed of one “aspirate” action followed by one “dispense” action. No “pull” actions occurred in the graphs. Pink shading in the Top and Middle graphs represents the summed feeding duration, while the Bottom graph breaks down the specific pump actions performed. Additional actions triggered by feedback are outside the pink feeding window. **(d)** Firing rate dynamics in response to feeding events across ascending cycle conditions. Neural activity was analyzed using 90-second sliding windows (in 1-second steps) and normalized in two stages: first using contrast normalization (║;*x*║; = (*a*−*m*)*/*(*a*+*m*), where ║;*x*║; is the norm, *a* is the firing rate of a neural unit in each window and *m* is the mean firing rate across all windows for that unit within its recording) to account for individual unit firing rate differences, followed by z-score normalization at each time window against the no-feed control recordings to account for natural baseline firing variability. Z-scores were averaged across units within each feed volume condition (150 µL to 900 µL) to evaluate the influence of progressively larger feed volumes.

Figure 6c presents neural unit raster plots and the average neuron firing rate for representative recordings of each condition. Raster plots of neural unit firing over time show no change in unit activity during microfluidic manipulation (light purple overlay). The average firing rate did not show unusual variability during or after the feeding window and did not correlate to pump actions (Dispense, Aspirate, or Cycle). To further inspect this computationally, we performed normalization and z-score analysis (detailed description in Materials and Methods, Normalization and z-score calculation for effect of feeding during recording). Within each recording, a 90-second sliding window was applied to the spike raster with 1-second steps. The firing rate of a single unit at any particular window was normalized to the same unit’s average firing rate across all windows within a recording. This scales changes in activity for any neural unit to be comparable irrespective of average firing rate. For all non-feeding recordings (N = 28), we calculated the firing rate mean and standard deviation (STD) of all normalized units in each window. The results of the experimental conditions (Figure 6d) (N = 3 per condition) were calculated with z-scores generated for each neural unit in each window to relate how much firing rate changed with respect to baseline variability. The STD of ± 1 in relation to the non-feed activity are marked (dashed blue line). In the 10 minutes following the onset of microfluidic feeding, the largest increase in firing rate was +0.7 STD above the mean at 3.9 minutes (150 µL condition), largest decrease was 0.6 STD below the mean at 0.8 minutes (300 µL condition). Z-scores for all conditions (150 µL, 300 µL, 600 µL, and 900 µL) remained within ± 0.7 STD showing no significant change in activity during microfluidic manipulations.

Feed cycles of different total volume and time length did not elicit immediate changes in firing rates during or minutes after feeding. Combined with our findings in Figure 5 showing no changes in firing rate distributions for six hours following feeding cycles, these results suggest that neural activity remains stable despite media exchange and the associated fluid movement.

## Discussion

This automated platform advances orgnaoid research methodology, enabling continuous monitoring and manipulation of brain organoids while maintaining optimal cell culture conditions through non-disruptive, automated protocols. It addresses issues of manual handling, variability, and limited temporal resolution. By integrating microfluidics, electrophysiology, and imaging through an IoT framework, we’ve created a system that supports experimental reproducibility and reveals temporal dynamics previously difficult to capture in manual protocols. Running on a distributed IoT network offers dual benefits: a local MQTT broker ensures reliable performance even during internet outages while Cloud integration enables global collaboration across distant labs for shared or complementary research. This setup enhances the continuity of individual experiments and the integration of worldwide scientific efforts. The reduction of human intervention enabled by the microfluidic feeding system reduces the risk of contamination, variations in time outside the controlled temperature and CO2 incbator environment during feeding and imaging, and other human-introduced variance. This degree of control is particularly valuable in long-term organoid experiments towards reducing batch effects.

Automated feedback mechanisms provide essential experimental control by maintaining conditions within defined target ranges without manual supervision. Here, we demonstrated one method of feedback: computer vision to maintain a consistent volume in the organoid growth chamber. During our 7-day studies, the system achieved this feedback autonomously and did not require manual intervention to rectify anomalies. Further applications of feedback using existing hardware could modulate electrical stimulation, media variety, or frequency of feeding based on media collections, morphology assessments, and electrophysiological measurements. Devices can use the flexibility of MQTT messaging to allow for the creation of additional feedback loops to control the experiment. The computer vision techniques we applied to volume estimation could be extended to further applications such as colorimetric and absorbance sensing using the same setup to interrogate biochemical properties of the media. Such measurements could provide additional analysis of organoid cultures leading to a nuanced understanding of their behavior and responses to different stimuli.

The interval between electrophysiological recordings is essential for characterizing neural network dynamics. Neural processes unfold with remarkable complexity and variability, but for practical reasons, many experimental paradigms are limited to once-a-day recordings [5, 40–42]. Recent work [43] demonstrated that important neural network properties, including firing rate distributions and small-world topology, are “preconfigured” rather than emerging solely through experience-dependent processes. Their finding that stable network properties exist from very early developmental stages validates that automated maintenance and monitoring systems, like the one presented here, can reliably capture intrinsic developmental processes without disrupting natural network organization. By providing the ability to schedule recordings at any interval, our system is particularly well-suited to investigate the relationship between innate and experience-dependent aspects of network development. Our high-frequency recordings revealed trends not captured in once-a-day sampling, enabling the detection of patterns, oscillations, and interactions that may be overlooked in sporadic recordings [44, 45]. These capabilities are further relevant for studying phenomena on shorter timescales, such as neuroplasticity [46], circadian rhythms [44], and for investigating neurodevelopmental disorders hypothesized to be ’connectopathies,’ characterized by abnormal connectivity [47]. By enabling simultaneous tracking of morphological, electrophysiological, and network-level changes over extended time periods, our automated platform could help resolve questions about how early network properties evolve throughout development, potentially yielding new insights into both the stability and plasticity of developing neural circuits.

The greater the complexity of experiments, the more automation becomes essential to coordinate and manage the different technical modalities. The use of 3D printing technology enhances this flexibility, allowing for the seamless combination of multiple systems, such as the integration of our custom media exchange setup with the commercial HD-MEA and portable microscope. We foresee the integration of additional sensory data and feedback mechanisms to analyze cell culture conditions. The lack of effect due to media manipulation presented in Figure 6 opens the opportunity dispense and aspirate pharmacological reagents or small molecule factors without the perturbation of manual interventions. With this system one could precisely measure the onset of electrophysiological responses to chemical manipulation of the culture. The platform’s consistency and reliability are ideal for comparative studies involving organoids of different genotypes or subjected to pharmacological manipulations. This capacity to facilitate direct comparisons between diverse experimental conditions in controlled environments holds promise for advancing our understanding of neurodevelopment and neurodevelopmental disorders.

## Materials and Methods

### Assembled devices and custom 3D-printed components

The Bill of Materials listing components and costs are provided in Supplementary Materials. STL files for 3D printing are provided in PrintedAccessories.zip.

### Microfluidic cell culture

The automated microfluidic pump system builds on previous work [30]. The microfluidic system was configured for this study to support two chips (AF and AFAR) and their respective collection reservoirs (right and left) were imaged by the camera setup. Replicates of the conditions were achieved by repeating the experiment on a following batch of organodis.

Fresh cell culture media is kept at 4°C refrigeration and accessed by the pump through flexible FEP tubing routed into a benchtop refrigerator and to a media bottled with a reagent delivery cap (Cole-Parmer VapLock). Fresh media is kept refrigerated to increase longevity and may be replaced during experimentation. To dispense, the syringe pump and distribution valves draw fresh media into the syringe vial and distribute the programmed volume into flexible FEP tubing routed through an access port in the incubator. Here, the media is heated in incubator conditions prior to being delivered to the organoid inside the culture chamber. To keep media dispenses available on demand, a preheated 450 µL reserve (59% of the chamber’s volumetric capacity) of fresh media remains idle in the FEP tubing so that upon dispensing, 37°C media is delivered to the well in less than 10 seconds. The FEP tubing is interfaced with the fluidic module with threaded ferrule lock and nut fittings (Cole-Parmer VapLock). Outflow from the fluidic module is drawn away with FEP tubing routed out of the incubator and into a refrigerator containing the collection reservoirs and computer vision camera setup.

For the collection reservoirs, we selected 15 mL Polyethylene Terephthalate (PET) conical tubes (430055, Corning) for high optical clarity, ease of replacement, and durability in downstream analysis and cold storage. To enhance visibility for computer vision imaging, we removed the factory-printed writing area on the conical PET tubes using generic, multipurpose tape. Flexible FEP tubing was interfaced with the PET tubes using a rubber cork plug (#6448K95, McMaster-Carr). The cork was pierced with 8-gauge steel needles that served as supportive conduits for the tubing. The tubing was secured inside the needle with glue (Loctite 4011) to create a hermetic seal at the point of interface. The steel encasing of the needles ensures a smooth, unobstructed flow within the flexible FEP tubes. Each collection reservoir had two flexible FEP tubes: one for media coming from the fluidic module and one for pressurized operation connected to the syringe pump. This ensured that spent media never entered the syringe (only air). The air is expelled into a filtered (Millipore AA 0.22 µm syringe filter) safety container (not shown in Figure 1).

For the 7-day studies described here, we designed for equivalent media exchange across conditions. The Controls were fed 4 times at 1 mL per feed, totaling 4 mL of replacement media. AF and AFAR were fed 28 times at 143 µL per feed, totaling 4 mL of replacement media over the week. Summing the scheduled feeds and feedback adjustments, a single collection reservoir could store conditioned media for 2-3 weeks.

### Cell culture

Cell culture protocols and organoid plating on HD-MEA, which occurred prior to the experiment, are described in Supplementary Materials and Methods.

### Priming the experiment

Before starting a 7-day recording experiment, membrane lids for HD-MEAs (AF and AFAR) were replaced with microfluidic culture chambers. During the replacement process, all media was aspirated from the HD-MEA’s well with a P-1000 pipette. The microfluidic catch tray, followed by the culture chamber, was inserted inside the well, and 750 µL of the original media was added back to the microfluidic culture chamber. Excess media was discarded. The glass rod lid was placed on top.

Flexible FEP tubes (idling with DI water) were flushed with 1.0 mL of fresh media. After priming the lines with media, the AF/AFAR chips were connected with fluidic fittings wrapped with Teflon tape. An initial aspiration leveled the media to the target fluidic operating range. The collection reservoirs were replaced with new empty conical tubes.

#### Running the experiment

During the experiment, the media was exchanged using a feed cycle operation consisting of an aspiration followed by fresh media dispense. Here, we performed 143 µL aspirations and dispenses every 6 hours to match 1.0mL feeds every two days in the manual feeding controls. Feedback performed additional aspiration, dispense, and pull actions in addition to the basic feed cycle schedule to ensure the system stayed within normative error ranges. See section Feedback interpreter.

#### Teardown of the experiment

Once the experiment was stopped, chips were disconnected from the flexible FEP tubes by unscrewing the fittings. The flexible FEP tubes with fittings were sterilized in a flask containing disinfectant (Cydex) and covered with aluminum foil. The collection reservoirs with the experiment’s conditioned media were disconnected and taken for analysis. New collection reservoirs were inserted for the cleaning cycle. The pump ran a cleaning solution (Cydex) through the entire internal cavity for 1 hour to disinfect the system. Following disinfection, DI water and dry, sterile air were profused through the system for 12+ hours (overnight) to clear the disinfectant. The flexible FEP tubes were left resting with DI water until the next experiment.

### Computer vision for fluid volume estimation

The computer vision setup, located inside a 4°C refrigerator, included a support for the collection reservoir, a camera module, and an LED panel positioned behind the conical tubes. The LED panel served as backlighting to enhance the clarity and contrast of the images. The reservoir support was a two-plex 3D-printed system capable of multiplexity to tailor alternate experiments (see Assembled devices and custom 3D-printed components). The camera and LED panel were both controlled by a Raspberry Pi.

To generate the calibration dataset, the camera module captured images of media in the collection reservoirs at select volumes over the entire range of the tube (0-12 mL), totaling 184 images. The volumes associated with each image were measured using a high-precision scale (30029077, Mettler Toledo). This approach enabled a correlation between the visual representation of media in the images and its actual volume (see Results).

To ensure image quality, our study introduced two checks to validate the integrity of the captured images: Lighting and blurriness. A region of interest (ROI) was designated within the panel’s area to verify the lighting conditions by checking that the average RGB color values each exceeded a minimum threshold of 20 out of 255. Blurriness was assessed by computing the variance of the Laplacian for the image, with a necessary threshold of 50 to pass. The thresholds were empirically determined using the calibration dataset.

Figure 2c illustrates the methodology applied to fluid segmentation, outlined in the Results section. The process begins with capturing an RGB image of the collection reservoirs that are fixed in place by the setup. To facilitate better segmentation and feature extraction, the RGB image is transformed into the HSV (Hue, Saturation, and Value) color space. A summation of the HSV values row-wise from the bottom to the top of the collection reservoir results in three distinctive profiles that allow differentiation between the liquid and background. Each profile, as illustrated in Figure 2c, presents a vertex at the boundary. A row value was established by averaging three rows identified in each HSV channel: an abrupt rise in the curve for the Hue channel, the absolute maximum for the Saturation channel, and the absolute minimum for the Value channel. From the average row value, the first segmentation was created. Everything below this row was set as white pixels, and everything above it was set as black pixels. A local evaluation around the average row was made to incorporate the meniscus in this segmentation. Utilizing HSV thresholds, the meniscus was accurately characterized and incorporated into the initial segmentation, culminating in the final image segmentation, in which white pixels represented the liquid portion.

The estimated volume was given by Equation 1, where x represents the segmented area in pixels, and the resultant volume is in microliters. Two different curves are used to account for the conical section for volumes under 1.5 mL (and pixel area less than 4446) and the cylindrical section for larger volumes.

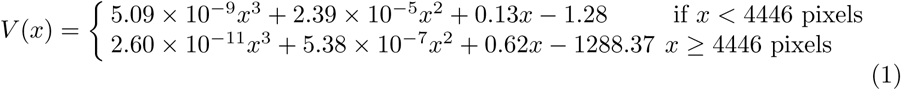

The image segmentation and estimation based on the mathematical model (Equation 1) is carried out by a software program named the “Estimator.” The process initiates with a feeding cycle, which triggers a picture request. Upon receiving the image of the collection reservoir, the “Estimator” analyzes the image and returns the estimated value of the fluid volume. The volume is relayed to the next module for feedback interpretation within the pump system (see Feedback interpreter).

### Feedback interpreter

Computer vision volume estimations were compared to expectation values based on the sum total of pump action jobs. The feedback interpreter classified estimations into four categories: within tolerance, out-of-tolerance, anomaly, and tube change. Tolerance was a static volume selected at the start of the experiment. For the results shown here, the tolerance was 143 µL. If the volume estimation received was within the expectation value +/- the tolerance, the pump action was determined a success, and feedback ceased. If the volume estimation received was beyond the expectation value +/- the tolerance and also less than +/- 2000 µL, another cycle of feedback was engaged. When the volume was less than expected, for the first 5 iterations of feedback, aspiration jobs were sent to the pump with the difference of expectation and estimation. For iterations 6 to 19, pull jobs were sent to the pump, increasing by one for each subsequent interaction. A “pull” is a 1000 µL aspiration at 10× the standard syringe speed (applying a 1.1 x 103 mm/s flow rate), shown to generate the force required to break through variably high resistance in the conditioned media. At 20 iterations, the feedback interpreter requests manual intervention via the messaging application, and all further pump actions are suspended until the issue is resolved.

When the volume was more than expected, dispense jobs were sent to the pump with the difference of expectation and estimation. Dispense actions were limited to 200 µL per action and 2 iterations of feedback in total to prevent overflow. A volume estimation that was 2000 µL or more above the expectation value was determined as an anomaly and requested manual intervention via the messaging application, and all further pump actions were suspended until the issue was resolved. The feedback interpreter automatically detected collection reservoir tube changes when the volume estimation dropped by 2000 µL or more compared to the previous estimation and the total volume present was estimated as less than 2000 µL.

### Computer vision for in-incubator organoid culture imaging In-incubator imaging

A 5MP digital microscope (AM7115MZTL, Dino-Lite) was placed over the organoid culture on the HD-MEA using holders described in Assembled devices and custom 3D printed components. Imaging was performed from the top through a glass rod (quartz drawn rod, 5mm ± 0.20mm dia x 15mm ± 0.20mm long, UQG Optics) (in AF/AFAR chips) or through a membrane lid (in control chips). The image is captured using reflected light from a built-in brightfield LED source next to the camera sensor. The 3D printed alignment trays handle most of the chip placement, with initial minor focal plane adjustment required. The microscope remains shut off until the software triggers it to turn on the lights and take a photo.

#### Image segmentation for organoid

In the process of image segmentation for organoid analysis, the first step involves applying an image calibration to correct any distortion. This procedure requires identifying four source points and four destination points. The former were manually selected from the distorted image. The latter were calculated based on an initial pixel (left corner of the HD-MEA), the size of the electrodes, and the spacing between them, both in millimeter units. This relationship between pixels and millimeters was established by using known dimensions of the HD-MEA border and electrode pitch in the image. The organoid segmentation within the rectified image was accomplished using the Segment Anything Model (SAM) [48]. This model combines neural network architectures, allowing for precise and versatile image segmentation without requiring specialized training on new images. The segmented image is analyzed to detect variations in pixel intensity, which signify the presence of organoid contours. Both images with the organoid’s contour and electrode grid are overlayed. Each electrode area is checked for the presence of the organoid’s border. When a border is detected within an electrode’s bounds, that particular electrode is marked prominently on the grid image to signify contact with the organoid (see Figure 5b). The step-by-step illustration of the analysis process is shown in Supplemental Figure 4.

#### Plotting & alignment to neural activity data

Electrode numbers as (x,y) position were plotted in matplotlib and exported as SVG. The SVG aligns over other plots, such as activity heatmaps, which follow the same x:3580 by y:2100 axis dimensions. Since electrophysiology plots use the electrode coordinate system with the same (x,y) positions, the image segmentation grid and neural activity plots are aligned on the same coordinate system.

### Measuring neural activity

Extracellular field potential recordings were performed using CMOS-based highdensity microelectrode arrays (HD-MEAs) (MaxOne, Maxwell Biosystems). Each HD-MEA contains 26,400 recording electrodes within a sensing area of 3.85 mm × 2.1 mm (each electrode has a diameter of 7.5 µm, spaced 17.5 µm apart center-to-center). A subset of up to 1020 electrodes (defined spatially by a configuration) can be selected for simultaneous recording [49]. Across one configuration, neuronal activity in microvolts was sampled over time at 20kHz and stored in HDF5 file format.

The experiment involved each chip’s daily activity scans and recordings (described below). Each chip underwent an activity scan and subsequent recording every day, consistently conducted within the same one-hour time window. All chips shared the same recording unit and were recorded one at a time. For the AFAR condition, beyond the daily recordings and activity scans, the chip remained on the HD-MEA for automated hourly recordings. The gain was set to 1024x with a 1 Hz high pass filter for both activity scans and recordings. The recording was set up to save 5 RMS thresholded spike times as well as all raw voltage data for downstream analysis and plotting.

All neural activity measurements were performed inside the incubator at 36.5°C, 5% CO2.

### Normalization and z-score calculation for effect of feeding during recording

In section Effects of feeding during recording, Figure 6d, the firing rate variability to feeding was derived as follows:

For every recording, a 90-second sliding window was used to further analyze the sorted and curated spike raster with steps of 1-second. For every window, the average firing rate per unit was computed. To account for intrinsic variability in firing rate between units, the average firing rate per window was normalized for each unit as ║;*x*║; = (*a* − *m*)*/*(*a* + *m*), where ║;*x*║; is the norm, *a* is the window rate and *m* is the mean value across all window rates in the recording. This normalization yields a values between -1 and 1, reflecting the deviation from the average firing rate of each unit. A value of 0 means that the average firing rate for a unit in a given window is the same as the average firing rate for that unit. A value of ± 1/3 means that the firing rate for a unit in a given window is approximately twice (for +1/3) or half (for -1/3) the mean firing rate for that unit.

Subsequently, the normalized firing rates per unit for all no-feed recordings were taken and for each window relative to the start of the recording, the mean and standard deviation were computed over all units in all no-feed recordings. These values were then used to z-score normalize the normalized firing rates of every unit in every window throughout all feed recordings as *z* = (║;*x_feed_*║;−*µ_nofeed_*)*/σ_nofeed_*, where *z* is the zscore, ║;*x_feed_*║; is the norm in the feeding condition, *µ_nofeed_* is the mean of the norm in the no feeding condition, and *σ_nofeed_* is the standard deviation of the norm in the no feeding condition. This normalization yields a value for each unit per fed recording at every window that reflects the variability from the average firing rate of that unit. This is expressed as the number of standard deviations that any particular value is removed from the average variability from the mean firing rate of all units over all no-feed recordings at that same window in the recording. If the firing rate of the unit during any point in the feed recording substantially increases or decreases relative to the average firing rate over the whole recording and if this is not the case for units in the no-feed recording, this will be reflected as a positive or negative z-score normalized value. In addition, the z-scores were averaged over all units in all feed recordings with the same feed volume to see the effects for each individual feed volume.

### Internet of Things (IoT)

#### Cloud Infrastructure

The cloud infrastructure, including S3, MQTT messaging, and cloud processing within the IoT system, has been previously described [28]. Additionally, we added a database service and defined a consistent organizational structure for MQTT messages and topics across devices and cloud jobs.

We use a combination of self-hosted services running on a server, and large data storage and analysis are performed on the National Research Platform (NRP) cloud compute cluster [50]. The devices are integrated with these cloud services:

- S3 cloud data storage: file storage using S3 object store, hosted on NRP cloud.
- Database: Strapi database stores device states, is self-hosted on our server, and is backed up to S3.
- MQTT messaging: EMQX MQTT broker, self-hosted on the server, and a Python messaging library (braingeneers.iot.broker) utilized by all software endpoints to send and receive messages from the broker.
- Cloud jobs/processing: utilizes a Kubernetes cluster on NRP and launches jobs. Employs software modularized by Docker containers and orchestrated by Kubernetes.
- User interfaces: features a website and integration with messaging apps (e.g., Slack) for interaction with devices, self-hosted on the server.

All custom software functionalities run in Docker containers and operate in a microservice architecture: specialized to a specific task and interface with minimal dependencies. A reverse proxy shields all web services from direct exposure to the internet. For example, webpages are configured through a reverse NGINX proxy, which not only assigns a specific domain to each service but also handles SSL and authentication services.

#### Security

Devices initiate communication with the server. Devices take MQTT commands in a specific format and are limited to the set of their defined commands, making them robust to command injection attacks. Accessing all cloud services requires authentication with credentials. All web, MQTT messages, database, and S3 storage operations are encrypted. Access to the user interface website is restricted through the proxy with a login authentication step. On the server side, all web-based microservices are secured through an NGINX proxy. The proxy allows web-based services to be relatively untrusted by providing security (https, authentication, internet visible network listener) and keeping all other web-based services on an internal docker network inaccessible from the internet. This simplifies security for services which may change often and accommodates programmers with minimal security training.

## Data availability

Electrophysiological data will be made available on a DANDI public server at: https://dandiarchive.org/dandiset/001268.

Other data including Bill of Materials, CAD models, and code will be available in the GitHub repository: https://github.com/braingeneers/integrated-system-v1-paper.

Additional data related to this paper may be requested from the authors.

## Acknowledgements

This work was supported by the Schmidt Futures Foundation SF 857 and the National Human Genome Research Institute under Award number 1RM1HG011543 (D.H., S.R.S and M.T.), the National Institute of Mental Health of the National Institutes of Health under Award Number R01MH120295 (S.R.S.), the National Institute of Mental Health grant 1U24MH132628 (M.A.M-R, D.H.), the National Institutes of Health (NIH) under award number K12GM139185 (J.L.S.) and the Institute for the Biology of Stem Cells (IBSC) at UC Santa Cruz, the National Science Foundation under award number NSF 2034037 (S.R.S and M.T), and NSF 2134955 (M.T., S.R.S and D.H). This work was supported in part by National Science Foundation (NSF) awards CNS-1730158, ACI-1540112, ACI-1541349, OAC-1826967, OAC-2112167, CNS-2100237, CNS-2120019, the University of California Office of the President, and the University of California San Diego’s California Institute for Telecommunications and Information Technology/Qualcomm Institute. Thanks to CENIC for the 100Gbps networks. The authors want to give special thanks to Jerry Glass, Anna Toledo, Ryan Hoffman, Demir Ozcakir, Quinton Brail, Viktor Yurevych, Kristof Tigyi, Samira Vera-Choqqueccota, Sugar Galka, Yohei Rosen, Valeska Victoria, Pierre Baudin, Rob Currie, Lon Blauvelt, Catharina Lindley, the IBSC Cell Culture Facility (RRID:SCR 021353), National Research Platform (NRP), and the UCSC Life Sciences Microscopy Center (RRID:SCR 021135) for valuable resources and assistance.

## Author contributions

- Conceptualization: KV, STS, MPM, DH, SRS, MT
- Data generation and curation: KV, STS, MPM, JLS, SH, HES
- Formal analysis: KV, STS, MPM, TM, DFP
- Funding acquisition: DH, SRS, JLS, MAM-R, MT
- Investigation: KV, STS, MPM
- Methodology: KV, STS, MPM, JG, TM, SH, HES, DFP, STM, TS
- Project administration: DH, SRS, MT
- Software: KV, STS, MPM, JG, TM, AR, DFP, DE, MATE
- Supervision: DH, MAM-R, SRS, MT
- Validation: KV, STS, MPM
- Visualization: KV, STS, MPM, JG, TM
- Writing – original draft: KV, STS, MPM, JLS, SRS, MT
- Writing – review & editing: all authors

## Supplementary Materials and Methods

### Embryonic stem cell culture

All experiments were performed in the adapted C57/BL6 mouse embryonic stem cell (ESC) line (Millipore Sigma # SF-CMTI-2). This line is derived from a male of the C57/BL6J mouse strain. Mycoplasma testing confirmed lack of contamination.

ESCs were maintained on Recombinant Human Protein Vitronectin (Thermo Fisher Scientific # A14700) coated plates using mESC maintenance media containing Glasgow Minimum Essential Medium (Thermo Fisher Scientific # 11710035), Embryonic Stem Cell-Qualified Fetal Bovine Serum (Thermo Fisher Scientific # 10439001), 0.1 mM MEM Non-Essential Amino Acids (Thermo Fisher Scientific # 11140050), 1 mM Sodium Pyruvate (Millipore Sigma # S8636), 2 mM Glutamax supplement (Thermo Fisher Scientific # 35050061), 0.1 mM 2-Mercaptoethanol (Millipore Sigma # M3148), and 0.05 mg/ml Primocin (Invitrogen # ant-pm-05). mESC maintenance media was supplemented with 1,000 units/mL of Recombinant Mouse Leukemia Inhibitory Factor (Millipore Sigma # ESG1107). Media was changed daily.

Vitronectin coating was incubated for 15 min at a concentration of 0.5 µg/mL dissolved in 1X Phosphate-buffered saline (PBS) pH 7.4 (Thermo Fisher Scientific # 70011044). Dissociation and cell passages were done using ReLeSR passaging reagent (Stem Cell Technologies # 05872) according to the manufacturer’s instructions. Cell freezing was done in mFreSR cryopreservation medium (Stem Cell Technologies # 05855) according to the manufacturer’s instructions.

### Cerebral cortex organoids generation

Mouse cortical organoids were grown as previously described by our group [34, 51] with some modifications. To generate cortical organoids we single cell dissociated ESCs using TrypLE Express Enzyme (ThermoFisher Scientific #12604021) for 5 minutes at 37°C and re-aggregated in lipidure-coated 96-well V-bottom plates at a density of 3,000 cells per aggregate, in 150 µL of mESC maintenance media supplemented with Rho Kinase Inhibitor (Y-27632, 10 µM, Tocris # 1254) and 1,000 units/mL of Recombinant Mouse Leukemia Inhibitory Factor (Millipore Sigma # ESG1107) (Day -1).

After one day (Day 0), we replaced the medium with cortical differentiation medium containing Glasgow Minimum Essential Medium (Thermo Fisher Scientific # 11710035), 10% Knockout Serum Replacement (Thermo Fisher Scientific # 10828028), 0.1 mM MEM Non-Essential Amino Acids (Thermo Fisher Scientific # 11140050), 1 mM Sodium Pyruvate (Millipore Sigma # S8636), 2 mM Glutamax supplement (Thermo Fisher Scientific # 35050061) 0.1 mM 2-Mercaptoethanol (Millipore Sigma # M3148) and 0.05 mg/ml Primocin (Invitrogen # ant-pm-05). Cortical differentiation medium was supplemented with Rho Kinase Inhibitor (Y-27632, 20 µM # 1254), WNT inhibitor (IWR1-*ε*, 3 µM, Cayman Chemical # 13659) and TGF-Beta inhibitor (SB431542, Tocris # 1614, 5 µM, days 0-7). Media was changed daily.

On day 5, organoids were transferred to ultra-low adhesion plates (Millipore Sigma # CLS3471) where media was aspirated and replaced with fresh neuronal differentiation media. The plate with organoids was put on an orbital shaker at 60 revolutions per minute. Neuronal differentiation medium contained Dulbecco’s Modified Eagle Medium: Nutrient Mixture F-12 with GlutaMAX supplement (Thermo Fisher Scientific # 10565018), 1X N-2 Supplement (Thermo Fisher Scientific # 17502048), 1X Chemically Defined Lipid Concentrate (Thermo Fisher Scientific # 11905031) and 0.05 mg/ml Primocin (Invitrogen # ant-pm-05). Organoids were grown under 5% CO2 conditions. The medium was changed every 2-3 days.

On day 14 and onward, we transferred the organoids to neuronal maturation media containing BrainPhys Neuronal Medium (Stem Cell Technologies # 05790), 1X N-2 Supplement, 1X Chemically Defined Lipid Concentrate (Thermo Fisher Scientific # 11905031), 1X B-27 Supplement (Thermo Fisher Scientific # 17504044), 0.05 mg/ml Primocin (Invitrogen # ant-pm-05) and 0.5% v/v Matrigel Growth Factor Reduced (GFR) Basement Membrane Matrix, LDEV-free.

### Organoid plating on microelectrode array

Mouse cerebral cortex organoids were plated, as previously described by our group [34], with two organoids per well. We plated the organoids at day 32 on MaxOne high-density microelectrode arrays (Maxwell Biosystems # PSM). Prior to organoid plating, the microelectrode arrays were coated in 2 steps: First, they were coated with 0.01% Poly-L-ornithine (Millipore Sigma # P4957) at 36.5°C overnight. Then, the microelectrode arrays were washed 3 times with PBS and coated with a solution of 5 µg/ml mouse Laminin (Fisher Scientific # CB40232) and 5 µg/ml human Fibronectin (Fisher Scientific # CB40008) prepared in PBS, at 36.5°C overnight.

After coating, we placed the organoids on the microelectrode arrays and removed excess media. The organoids were incubated at 36.5°C for 20 minutes to promote attachment. We then added prewarmed neuronal maturation media (described in the section above). We exchanged 1.0 mL of conditioned media for fresh every 2 days.

HD-MEAs containing the organoid cultures are stored in an incubator at 36.5 °C, 5% CO2, covered with membrane lids described in the section below, Assembled devices and custom 3D-printed components.

### Computer vision for fluid level detection

#### Camera details

A 16MP camera (B0290, Arducam) and a set of conical tubes are fixed 12 mm apart from each other on an optical breadboard (SAB10×30-M, Base Lab Tools). The camera was specifically configured without autofocus, with its focus statically set at 344 on a scale from 1 to 1023. A two-second warm-up period stabilizes the focus setting before a picture is taken. Exposure was set at 45 on a scale from 1 to 5000.

#### LED panel details

A 16×16 LED matrix (WS2812B-16×16ECO, BTF-LIGHTING) covered with 0.1mm thick polyester diffusion film (B08PTCGTX9, RENIAN) creates a uniformly illuminated background (we used 8 sheets of diffuser film spaced 1 mm apart by double-sided foam mounting tape). The LED panel is approximately 5 mm behind the conical tubes. The LED matrix was set to display a color gradient to best contrast fluid contents inside the conical tube, particularly in the cone-shaped lower area of the conical tube, which is thinner and appears lighter in color. The red color component of each LED matrix pixel was set based on its row position within the matrix, beginning with an initial red value of 221 out of 255. The red color intensity was reduced by two units for each row upwards, creating a gradient effect. Thus, the final color of each pixel was a combination of this dynamically adjusted red value and fixed green and blue values of 140 and 180, respectively. Furthermore, the LED panel’s brightness was set to 50% to prevent overexposure in the captured images.

### Assembled devices and custom 3D-printed components

All custom accessories were 3D printed (Form 3B+, Formlabs) with Biomed Clear V1 material (RS-F2-BMCL-01, Formlabs), except for the collection tube and camera stand in the refrigerator printed in BioMed Black V1 (RS-F2-BMBL-01, Formlabs). The parts were printed flat on the build plate to reduce support material. Alignment grooves between the insert and lid described in the Microfluidic culture chamber form a hole which also facilitates 3D printing by removing the formation of suction cups to the resin tank.

#### Microfluidic culture chamber

The microfluidic culture chamber assembly allows media to be exchanged inside the HD-MEA well. The chamber assembly consists of a microfluidic module, glass rod lid, and catch tray (Figure 1b,f,g).

The microfluidic module is placed inside the HD-MEA well, creating a media chamber and fluid path into and out of the chamber. Media from outside the incubator travels to the fluidic insert along 0.030” ID and 0.090” OD Tygon tubing (AAD02119-CP, Cole Parmer); the length of the tubing is approximately 100 cm. The tubing attaches to the fluidic insert using PEEK fittings (EW-02014-97, Cole Parmer) wrapped (counter-clockwise) in PTFE thread seal tape around twice the fitting’s circumference. The inlet and outlet are raised inside the fluidic insert to create a pool following a geometry published in previous work (30).

The fluidic insert, glass rod lid, and catch tray use silicone O-rings (5233T543, 5233T479, 5233T297, and 5233T585, McMaster) to provide seals against contaminations and leakage. O-rings were rubbed with a minimal quantity of canola oil for lubrication to facilitate installation and enhance sealing performance. The canola oil can be autoclave-sterilized, but it is unnecessary if the O-rings are sterilized post-lubrication (see section, Sterilization and assembly).

#### Membrane lid

The membrane lid used for experimental control conditions follows established designs (49), with adjusted dimensions to improve grip, matching material to the microfluidic culture chamber, and high-temperature silicone O-rings instead of rubber. The outer O-ring (5233T683, McMaster) holds the breathable FEP film (23-1FEP-2-50, CS Hyde Company) stretched over the top of the lid. The inner O-ring (5233T585, McMaster) seals the lid and well. The inner O-ring is also rubbed with a minimal quantity of canola oil as described in the Microfluidic culture chamber section.

#### In-incubator imaging alignment holders

The custom alignment holders, designed for two configurations, center a digital microscope over the biological sample on the HD-MEA. Components are screw mounted (91292A134, McMaster) to optical breadboards (SAB10X15-M, SAB15X15-M, Base Lab Tools Inc.) to ensure stability and maintain accurate spacing.

#### HD-MEA off the recording unit

The microscope is held over a single HD-MEA by a post and clamp (MS08B, Dino-Lite) mounted with a setscrew and base (SS6MS10, TH15/M, Thorlabs). The custom HD-MEA holder centers it for imaging. Throughout the experiment, HD-MEAs were left resting on each holder. The holder has cut-outs for handling the chip and also avoids the chip’s contact pads to decrease scratching and avoid moist surfaces. The holder also has indicators for the chip’s proper rotation with respect to the microscope.

#### HD-MEA on the recording recording unit

The custom holder on a post assembly (SS6MS10, TH15/M, TR250/M-JP, Thorlabs) mounts the microscope over the chip on the recording unit. The custom holder centers both the recording unit with its attached chip to the microscope.

#### Sterilization and assembly

Before use in tissue culture, components were placed in autoclavable bags (RIT-3565, PlastCare USA) and steam-sterilized at 134°C for 20 minutes or 121°C for 30 minutes based on Formlabs material datasheet specifications. Components were autoclaved, disassembled, and then assembled in a sterile tissue culture hood to avoid deformation or cracking during temperature cycling. Components were transported in an enclosed petri dish (small items) or a sterile autoclaved bag (large items) before being released into the incubator. Components that could not be autoclaved (such as electronics, i.e., recording unit, microscope) have their enclosures sterilized with hydrogen peroxide disinfecting wipes (100850922, Diversey) before entering the incubator.

### Measuring neural activity

#### Activity scans

Activity scans were performed daily in the MaxLab Live Scope (Version 22.2.22, MaxWell Biosystems) to identify where the organoid’s electrical activity is spatially distributed across the HD-MEA. The activity scan sequentially records from different configurations of up to 1020 electrodes, thereby sampling the microelectrode array for action potentials. We used the checkerboard assay consisting of 14 configurations, with 30 seconds of recording per configuration. The resulting activity heatmap (see Activity heatmaps) for each chip is shown in Figure 5b. Based on the assay results, 1020 most active electrodes were selected for simultaneous activity recordings.

#### Recordings

Each recording lasted 10 minutes. Initial recording configurations were created on the first day, and configurations were updated on the second day to match shifting activity. Afterward, we chose to keep the configurations constant across the final 5 days since the activity did not shift dramatically, and keeping the same configuration allowed for more consistent monitoring of the same region.

#### Smartplugs

A smartplug was connected to the recording system to automatically manage the duration of the recording system running. The smartplug (S31, SONOFF) running Tasmota 13.2.0 was connected to the MQTT broker (see MQTT) and received MQTT commands over WiFi to turn on and off.

The smartplug facilitated the automated recordings every hour: on the computer connected to the MEA recording system, a script running in Python (3.10) triggered the smartplug via MQTT to turn on the recording system, performed a recording using MaxLab Python API (MaxWell Biosystems), and afterward triggered the smartplug to turn off the recording system.

#### Spike sorting and curation

To process the electrophysiology data, each MaxWell recording was spike sorted into single unit activity using Kilosort2 [39]. Using a template-matching algorithm, Kilosort2 clustered neurons based on waveform shape. Spike sorting parameters included a bandpass filter of 300 to 6000 Hz for the raw data and voltage threshold of 6 RMS above baseline.

The sorting output was curated by an automatic algorithm that quality checks signal-to-noise ratio (SNR), firing rate, interspike interval (ISI) violation, and spike footprint for each putative neuronal unit. As a result, units that had SNR above 3, firing rate above 0.1 Hz, ISI violation below 0.5 and footprint on more than one channel were kept for analysis [52]. Units were labeled redundant using spikeinterface “remove redundant” module and processed through manual curation for consolidation.

Spike sorting was performed on the National Research Platform (NRP) computing cluster with an NVIDIA A10 GPU.

#### Activity heatmaps

Activity heatmaps in Figure 5a depict the spatial distribution of significant voltage events. MaxWell software provides thresholded event identification based on moving root-mean-square (rms) value for each electrode, identifying events exceeding 5 times an electrode’s rms value. We created a 2D grid of spike counts per second and applied a 2D Gaussian blur for visual smoothness, normalizing each grid point by dividing it by 2*πr*^2^ to re-scale back to the original Hz values. These values were then plotted as the activity heat maps. The heatmaps use warmer colors for higher firing frequency and darker colors for lower activity.

#### MQTT

MQTT messages serve as the standard unit of communication (Figure 3b, orange). MQTT allows devices and services to communicate without direct dependencies between each other by using a common publish/subscribe medium. MQTT clients are the devices or software entities that connect to the broker to send (publish) or receive (subscribe to) messages. Devices and services send messages on MQTT topics, which are hierarchical strings that allow listeners to capture a wide or narrow scope of information. Messages contain a payload with a list of key-value pairs to structure information. For example, a message requesting a microelectrode array to record has a key for recording duration with a value in minutes. Examples of MQTT topic structure and message JSON payloads are summarized in Supplementary Table 1; see GitHub for more information^1^.

The MQTT broker is the central communication facilitator in the network and coordinates messages between clients. The MQTT broker receives all messages from the clients, filters these messages based on their topics, and then distributes them accordingly to other clients who have subscribed to those specific topics. This setup enables efficient message routing and ensures that messages reach the intended recipients without the senders needing to know the specific details of the recipients.

Clients can be sensors, actuators, applications, and services (like UIs or analysis), or any other devices capable of network communication. The organization is future-proof because MQTT allows the creation of new services and devices and uses information available without changing any services (logging, UI, dashboards, analysis of traffic, etc.). Furthermore, message bridges can be employed to convert MQTT messages to other messaging APIs such as text messaging, email, or work chat applications like Slack (see Messaging bridge).

#### IoT device-class

The primary function of a *device-class* involves listening for job requests, executing them, and saving the resulting data to the cloud. This data includes measurements (e.g., images, voltage recordings) and log entries detailing device actions (e.g., cell culture feeding events). By consolidating features, the *device-class* framework simplifies the creation of new devices and enables easy control, updating, and interoperability. The Python *device-class* provides standard features across all IoT devices:

- a state machine defining standard behavior (i.e., experiment workflow)
- structured framework for processing incoming request messages
- autonomous task scheduling, timing, and execution; the internal scheduler manages time
- conflicts of tasks or autonomously recurring jobs
- multi-tasking and responsiveness to user requests via threading
- built-in database operations (i.e., updating device state (shadow))
- communication via MQTT messaging (including alerts via Slack bridge)
- background data upload/download mechanisms, managing queueing and retry
- error handling mechanisms
- communicate and work with other devices in a fleet

A child of the parent *device-class* will inherit all basic functionality, and may add additional features. For instance, a camera *device-class* child performs all actions that a *device-class* can, plus it knows how to handle a request to take a picture.

Having a common parent class consolidates similar features for different devices and allows for easier updates because all devices use the same core code library. The *device-class* code is available within the Braingeneerspy Python package on GitHub^2^. For state machine states and request commands see Supplementary Tables 1 and 1.

Devices can work in a fleet. As each device has the same core software with complementary behaviors, they integrate seamlessly, similar to how uniform building blocks can easily snap together. Devices can ask each other to yield while they perform sensitive actions (Figure 3d). Similarly, devices can perform services for each other in a coordinated manner. For example, midway through a recording, a microelectrode array device could ask the pump to deliver a drug. Devices can perform rudimentary decision-making to simplify overarching management. Devices post status and information to an open MQTT topic, allowing services and devices to build on and interface with those devices without altering existing devices and services. Devices can use each other to make sure the experiment is on track across multiple modes of sensing, for example the pump using the eyes of the camera to ensure pumping succeeded.

#### Pre-experiment workflow

Figure 3a illustrates the state transitions of a generic device during operation. It begins in the SHUTDOWN state, moving to IDLE, where it waits for user setup verification. Post-setup, it transitions to PRIMED, ready for experimental involvement. In the READY state, the device listens for experiment-specific MQTT messages, ignoring external recruitment until released with an END message. Devices can communicate collectively via MQTT topics for coordinated actions. Transitioning to WAITING occurs upon receiving a pause command, halting job execution. The device moves to EXEC when starting a job, returning to READY upon completion. Data uploads are managed independently of state changes, ensuring continuity even during outages. Devices can exit an experiment at any stage, reverting to IDLE or SHUTDOWN, with data upload tasks resuming upon restart. Figure 3a describes a generic device (e.g. a scientific instrument) and how it transitions between states during operation. On device start, the device transitions from SHUTDOWN state to IDLE. In the IDLE state, the device is waiting for a user to verify or install physical prerequisites. The IDLE state ensures the user performs the necessary setup of their device to maintain safety and usability. For example, a pump may wait in IDLE state until a user checks and confirms that the pump is clean and proper reagent bottles are connected. On the other hand, a camera may not have any prerequisites and would immediately transition to the next state, PRIMED. In the PRIMED state, the device has all the prerequisites to perform its job and waits to be called into an experiment. Devices listen on their default device MQTT topic. Once it receives a correctly formatted ‘start’ MQTT message (see ‘START’ message in Supplementary Table 1), it can transition to READY.

#### Experimental workflow

When the device transitions to READY state, when it listens to an MQTT topic for the experiment. It will refuse requests to be recruited to other experiments until it is released from the current experiment by an END message (see END message in Table 1). This ensures other users don’t accidentally disturb or recruit an occupied device into a parallel experiment. Switching MQTT topics also ensures exclusivity in incoming messages. The experiment topic structure (see MQTT) allows devices to send a group message addressing all devices. For example, a device or user could send a message to roll-call all devices on the topic (see PING message in Table 1) or pause all devices while it performs a sensitive action (see PAUSE message in Table 1). Upon receiving a message to pause, the device transitions to WAITING state, where it does not perform any jobs.

Once a device returns to READY state, it can transition into EXEC state if it receives a job request or has a job request from its schedule. If the device is in WAITING or EXEC while receiving a job request, it will put the request on the schedule to be executed as soon as possible. During EXEC state, the device is actively executing a job request. Once the job finishes or is stopped (see STOP message in Table 1), the device transitions back to READY state. Any data produced is queued for upload, protected from internet outages by upload retries with exponential backoff. Uploads occur in the background, independent of device state. A device can begin EXEC on a new job immediately after queueing the previous data for upload. From any state, a device can be terminated from an experiment and return to the IDLE state. At any point in the experiment, if a device is gracefully requested to turn off, it performs a final transition to SHUTDOWN state before halting the program. The device keeps the upload queue saved on disk and will continue unfinished uploads upon restart.

#### Data uploading

Data is saved to a ‘diskcache’ in memory. Once a file is produced, it is put on the upload queue. The upload queue contains references to files within diskcahe. Typical devices have at least 32 GB of disk memory, far larger than a single file. The queue is restricted to grow up to 80% of the device’s memory. Once the memory of the device fills up, older files that were uploaded can become overwritten.

#### Messaging bridge

The messaging bridge serves as an intermediary for communication between different platforms. It is a service that listens to MQTT messages in the IoT environment and translates them into other APIs like Slack.

The Slack bridge allows IoT devices to send notifications to individuals in designated Slack channels. The messaging bridge uses the message broker API and Slack API [53]. The Slack API requires an API key to be registered with Slack and an API bot to be added to the Slack channels of interest. The message bridge listens to an MQTT channel dedicated to Slack messages. When devices want to post a message to Slack, they publish a message on the dedicated Slack MQTT topic with a JSON payload containing the message. The payload can include text and image data. To support images, a link to an S3 object can be passed in the message, and the messaging bridge will then download and attach it to the Slack message. An image can also be sent directly inside the MQTT message, this requires modifying the message broker service’s configuration to increase the MQTT message buffer size to accommodate larger KB-sized files. The Slack bridge is a relatively simple service that decouples devices from dependencies on a specific API by communicating using the common message format MQTT.

#### Website

The website’s front end is developed using React, a JavaScript library for building dynamic and responsive user interfaces. For the backend, Flask, a lightweight Python web framework, is employed. Flask’s simplicity and flexibility make it ideal for our web services. It handles server-side operations, data processing, and interaction with databases.

The system’s structure incorporates a message broker API, which is established on the backend side of the architecture. This message broker is responsible for the asynchronous communication and management of all IoT devices connected to the cloud. Additionally, Flask’s compatibility with Python enables seamless integration with Python APIs, including the braingeneerspy MQTT message broker.

Through the front end, users can issue commands to the devices, and the message broker API in the backend efficiently manages these requests. The user interface encompasses three main components: the initialization page for entering initial experiment data, the control page for managing devices and monitoring their status, and the visualization page for analyzing experimental data through various graphs. All three pages require a specified experiment UUID (see Figure 3).

Both frontend and backend components are containerized using Docker, ensuring consistency and isolation in different environments. Integration of Cross-Origin Resource Sharing (CORS) is crucial for allowing the React frontend to securely interact with the Flask backend hosted on a different domain.

### Initialization page

On the initialization page, users can enter metadata containing experiment and biological sample details, which are compiled into a JSON file and uploaded to cloud storage, serving as a centralized repository for all experimental data.

### Control page

On the control page, users can access all the devices involved in the experiment associated with a specific UUID. For each device, users can request the execution of all the commands listed in Table 1, such as starting, stopping, and pausing the device, as well as scheduling tasks. Additionally, on the control page, users can monitor the real-time status of the device, as outlined in Table 1.

### Visualization page

On the visualization page, users can load data related to the volume estimator from current or previous experiments of a specific UUID. It is also possible to download images on a specific timestamp, allowing for manual monitoring of reservoir tubes.

**Supplementary Fig. 1.**
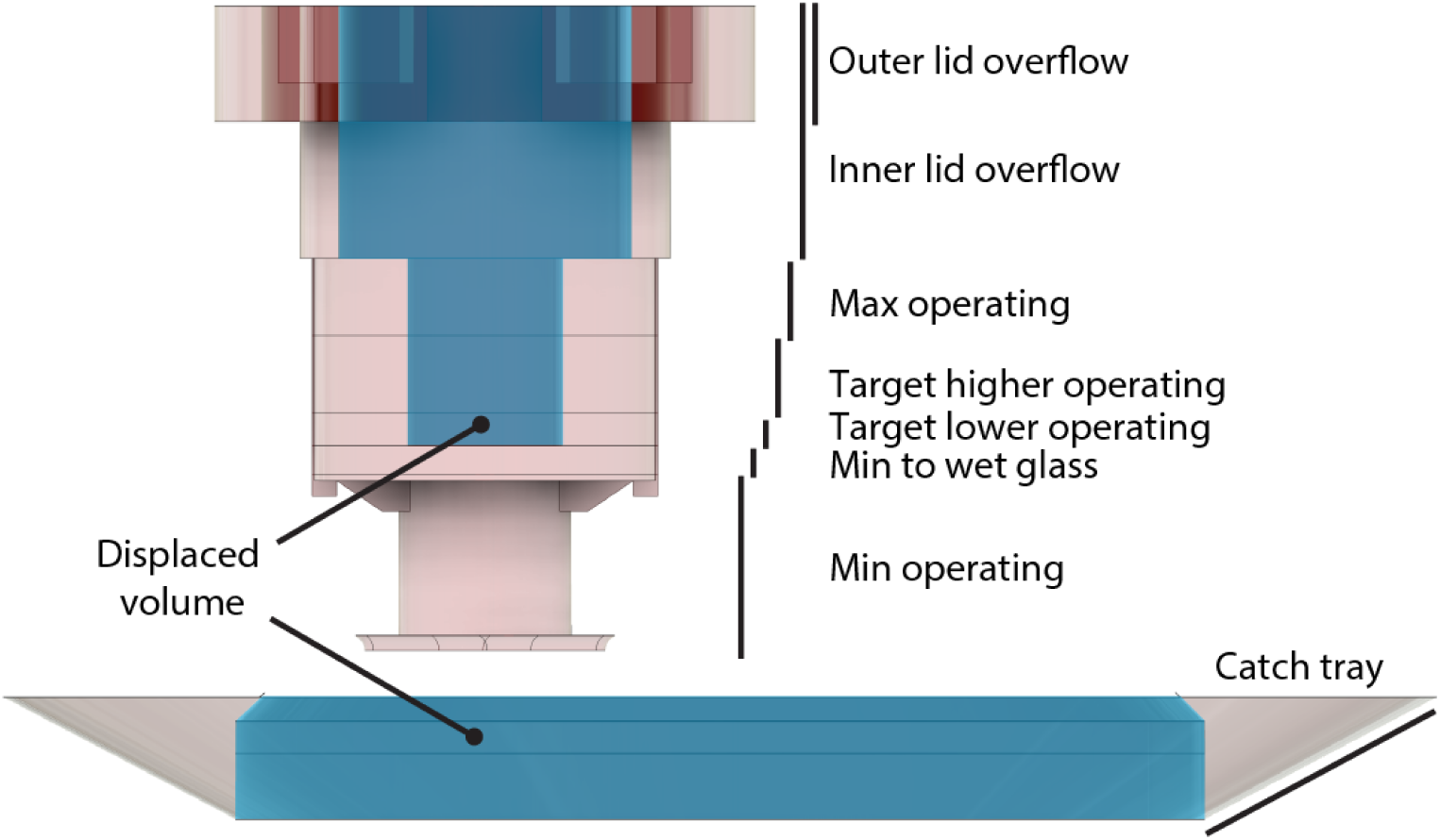
Diagram of operating ranges of the microfluidic culture chamber. Shaded pink areas represent volumes where media is collected. Shaded blue areas mark displaced volumes (where there is no media stored). The numerical volumes for each operating range are listed in Supplementary Table 1.

**Supplementary Table. 1.**
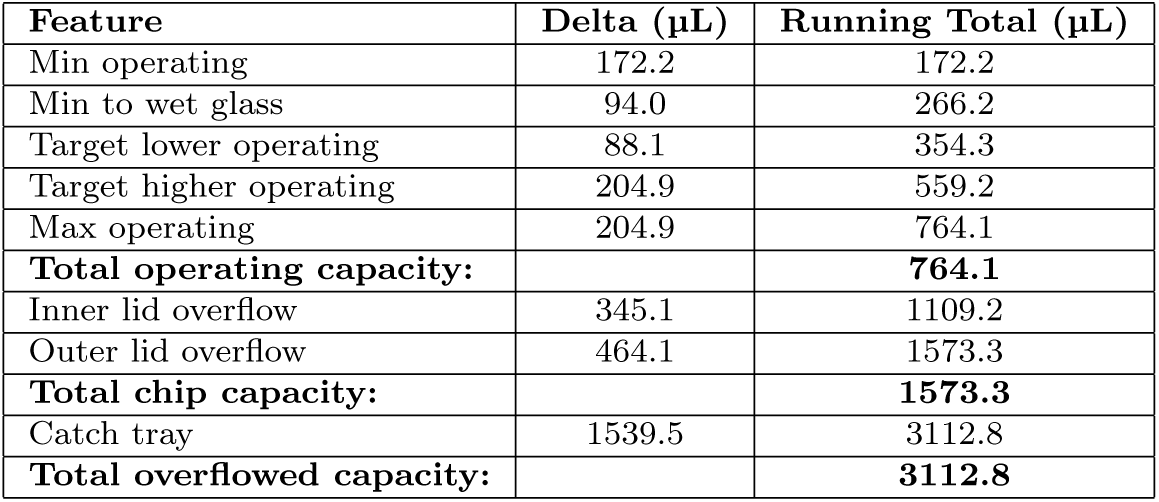
Numerical operating volume ranges based on the microfluidic culture chamber’s 3D model (CAD) measurements. Illustrations of operating ranges are shown in Supplementary Figure 1. The Feature column lists critical points in the microfluidic culture chamber. The Delta column is the volume space between each feature, and the Running Total column is the volume from the floor to the feature.

**Supplementary Table. 1.**
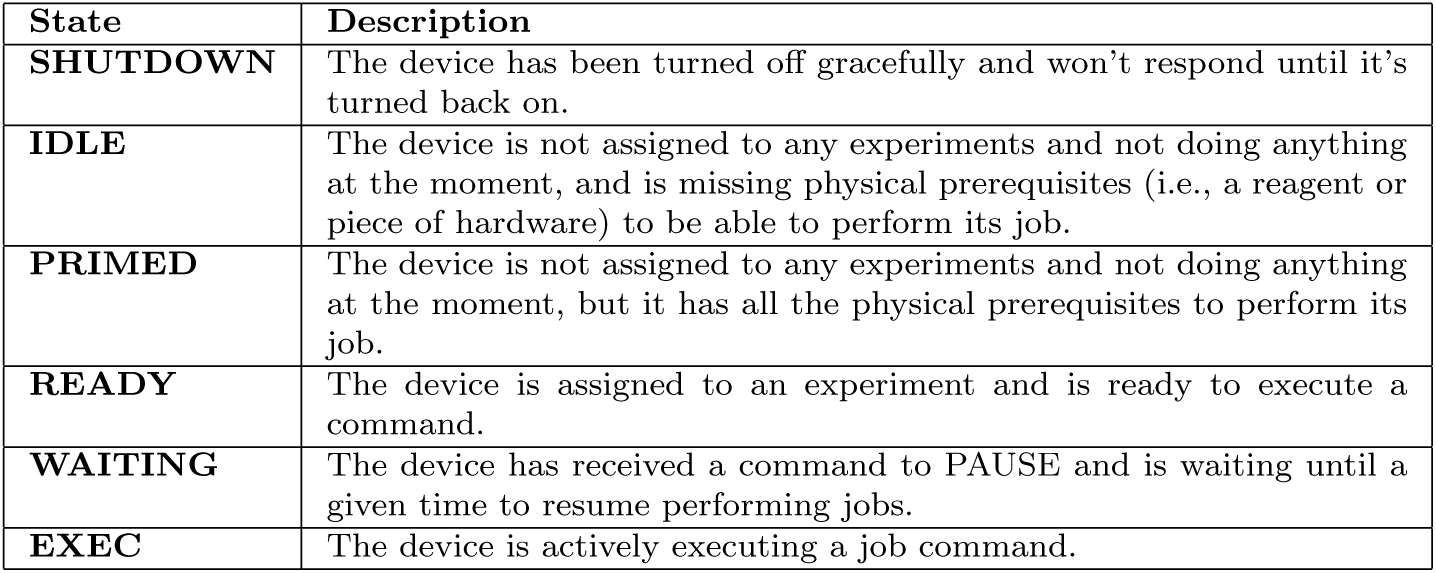
Device states. The *device-class* is structured as a finite-state machine, with a defined set of states (SHUTDOWN, IDLE, PRIMED, READY, PAUSED, EXEC) that describe its status. The finite-state machine reads a set of inputs and changes to a different state based on those inputs. The inputs can be user physical interactions (i.e., button press, linkage of consumables, etc.), MQTT messages containing job requests, or scheduled events.

**Supplementary Fig. 2.**
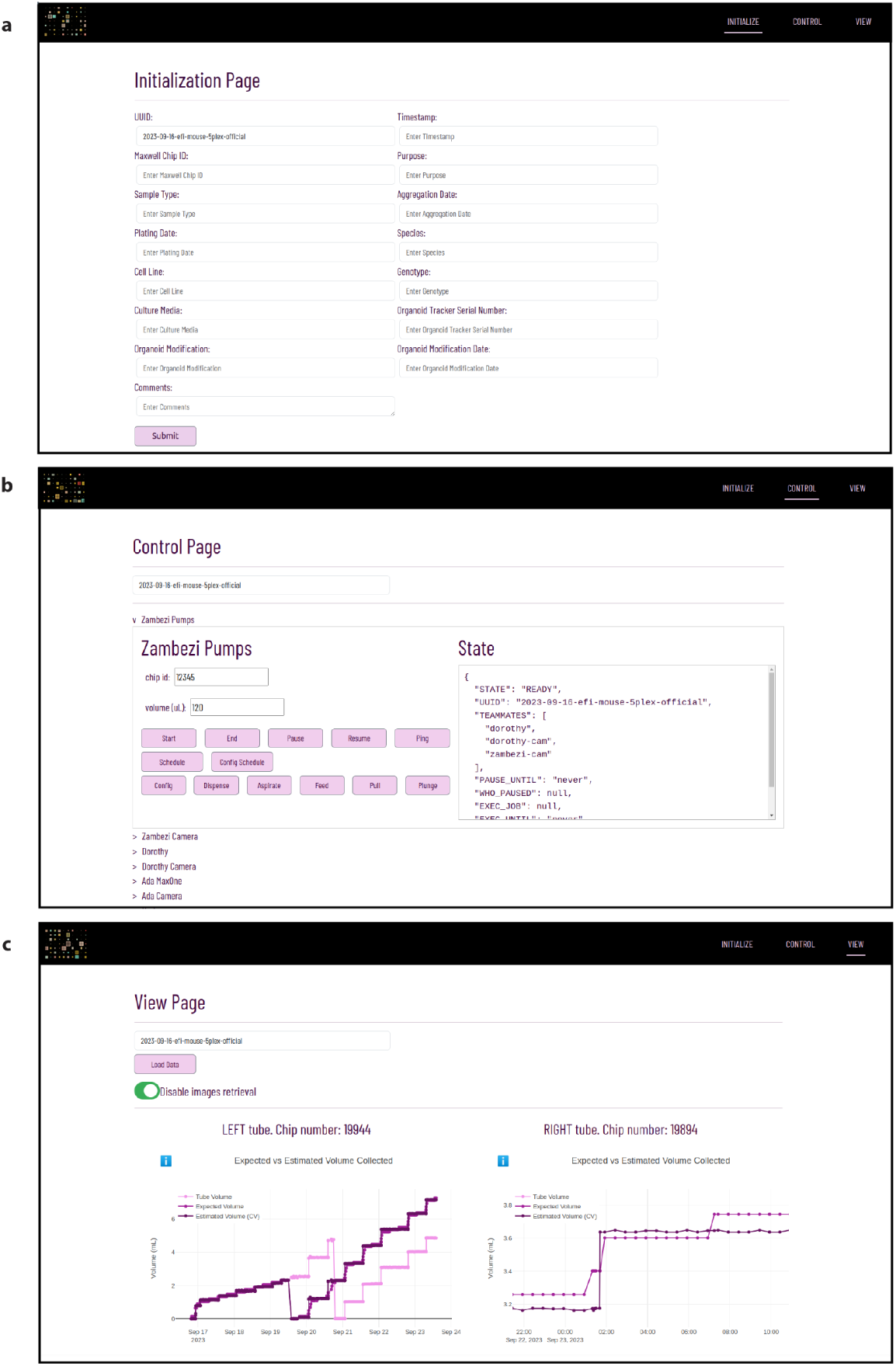
Webpage user interface screenshots. **(a)** Initialization page: Users can input details about the experiment and the biological samples. **(b)** Control page: Users can access and control every device involved in the experiment. **(c)** Visualization page: It includes three graph types. (1) Expected versus Estimated Volume Graph: compare volumes determined by the computer vision algorithm with volume according to pump metrics, highlighting any discrepancies and mismatching data. (2) Expected minus Estimated Graph: It shows the difference between the pump metrics and computer vision estimates for each device. They are designed to quickly identify alignment or discrepancies between these two methods, where values close to zero suggest good alignment, and deviations indicate measurement inaccuracies. (3) Collected Volume According to Computer Vision and Pump Graph: This graph contrasts the volume of media collected as reported by the pump system with that detected by the Computer Vision algorithm, which is crucial for assessing feeding accuracy. For example, if the pump indicates a feed of 300 µL, but the Computer Vision only detects 150 µL, this discrepancy is highlighted.

**Supplementary Fig. 3.**
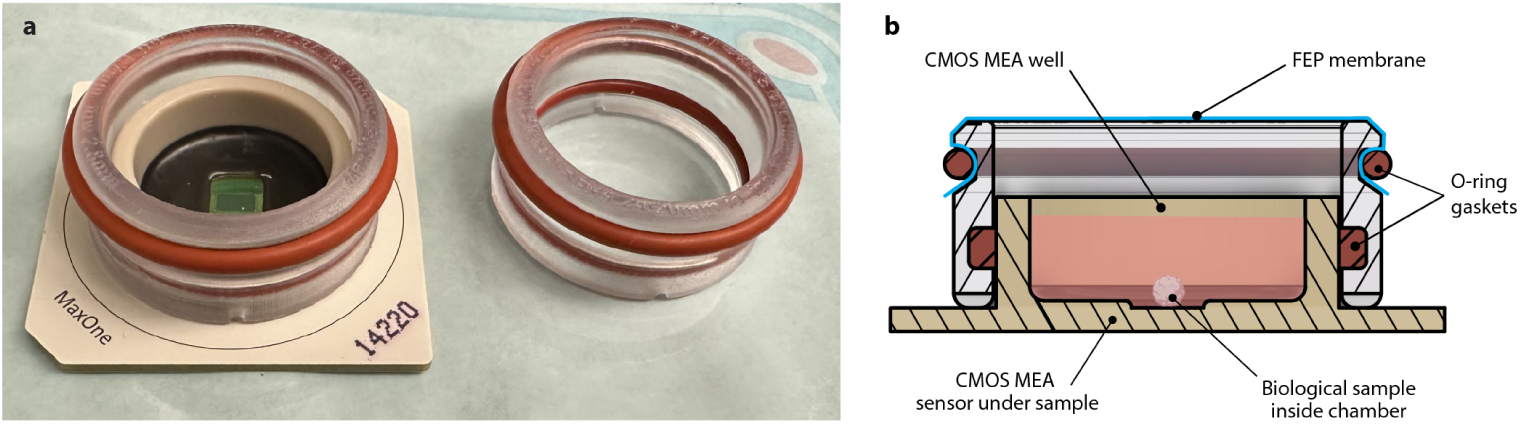
3D printed breathable membrane lid used for Controls modeled after designs by Potter. [54]**. (a)** Picture of the membrane lid and HD-MEA. The chamber is comprised of biocompatible 3D-printed parts, sealed by O-rings to the HD-MEA, and imaged through the FEP membrane stretched over the top with an O-ring. **(b)** Cross-sectional rendering depicting the fluid path and position of the sample.

**Supplementary Fig. 4.**
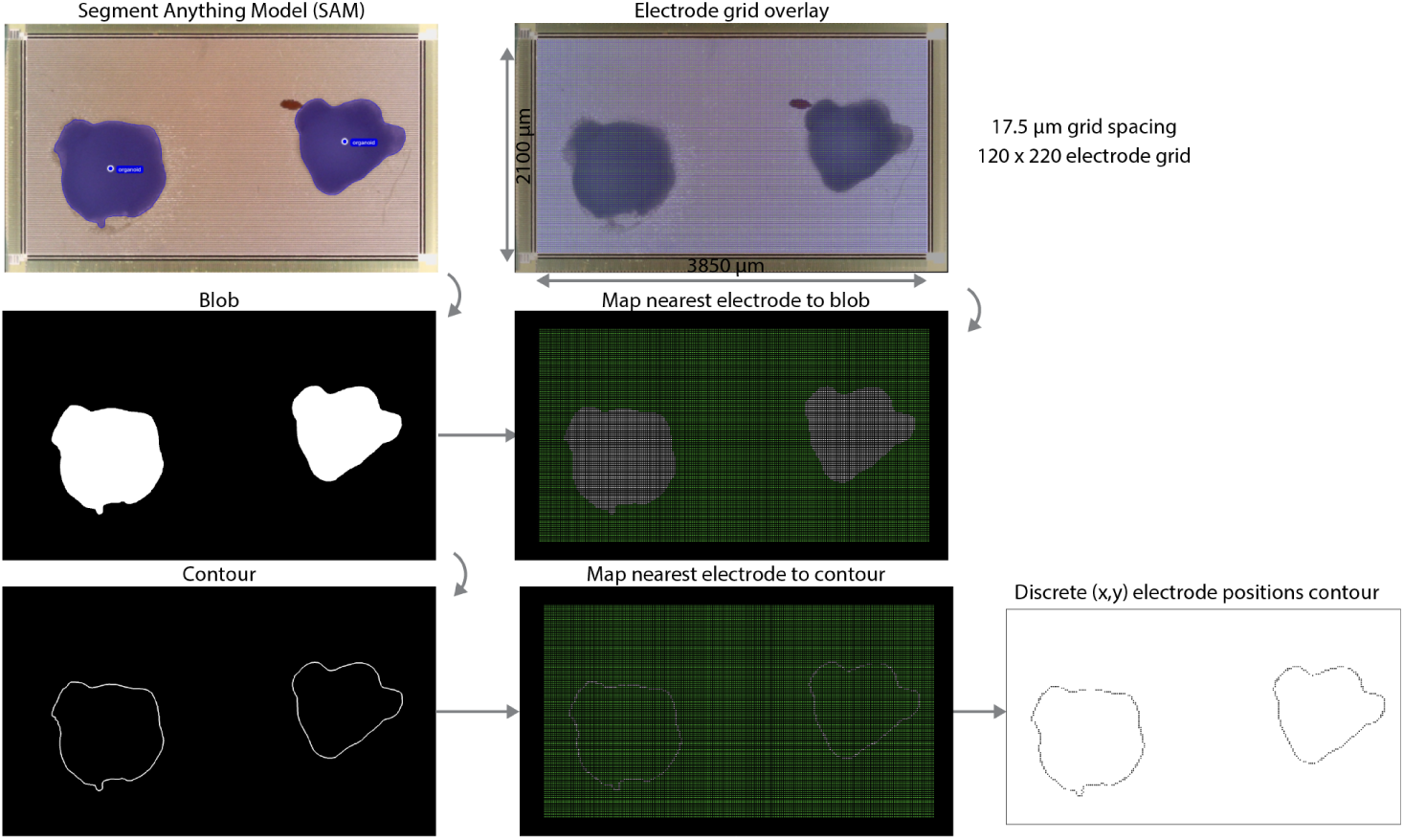
Organoid boundary segmentation process.

**Supplementary Table. 1.**
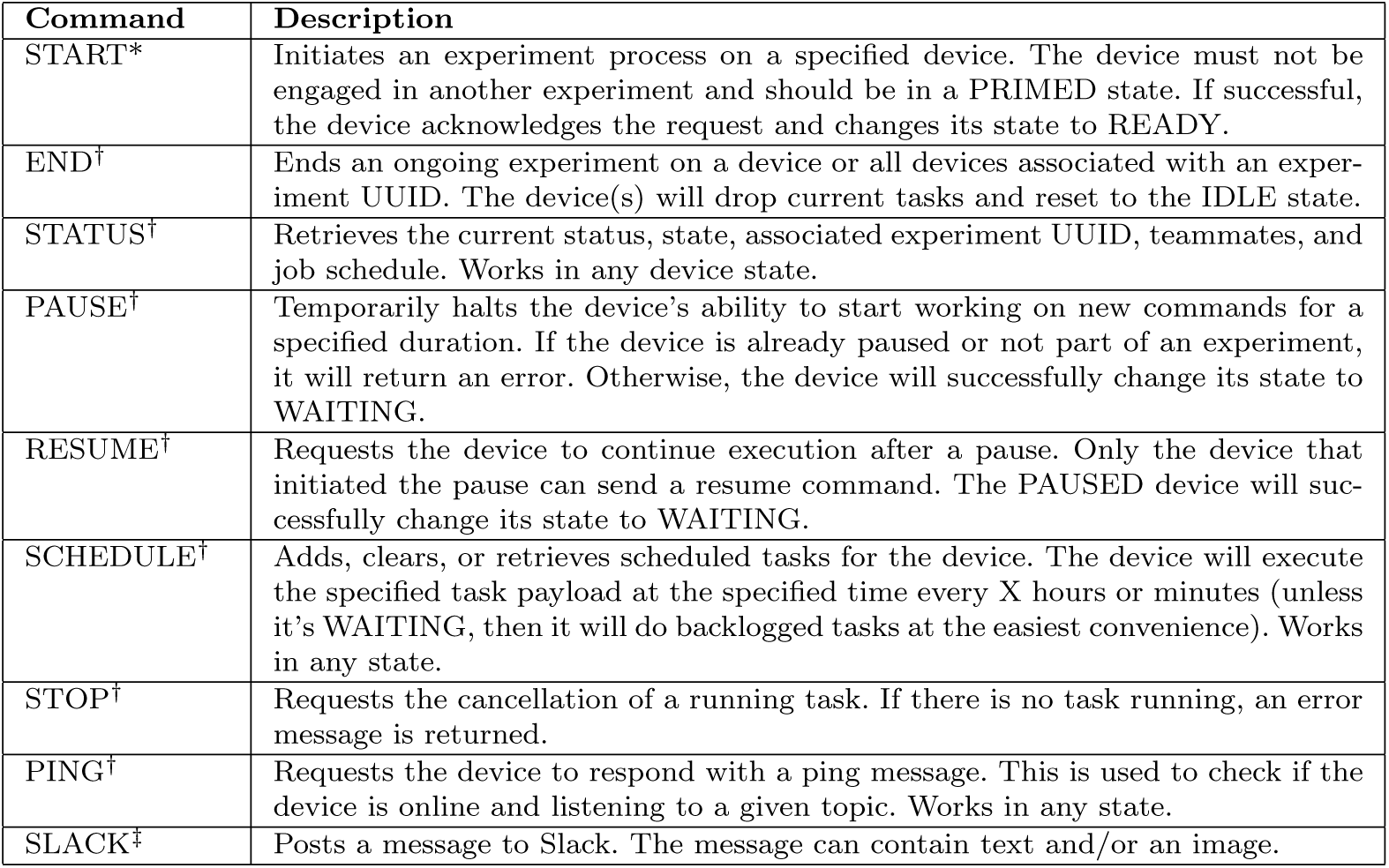
Generic commands. The parent *device-class* responds to a generic set of commands. Commands are sent on hierarchical MQTT topics that allow widening and narrowing of scope. We used each experiment’s Universal Unique Identifier (UUID) and each device’s name as part of the topic. If a device is not part of an experiment, the default UUID is NONE. * Use MQTT topic: NONE/device because no experiment assigned yet *^†^* Use MQTT topic: UUID/device or just UUID to address all *^‡^* Use MQTT topic: TOSLACK

**Supplementary Table. 1.**
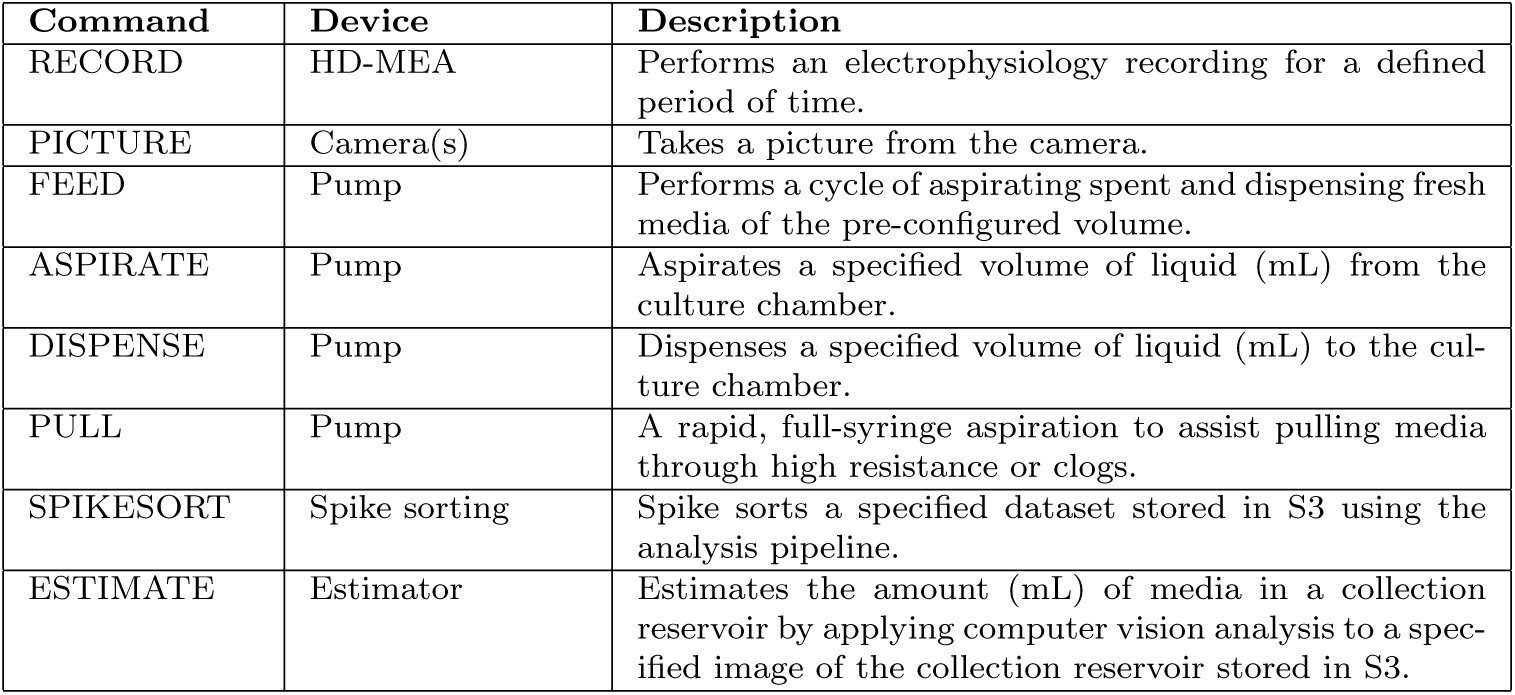
Application-specific commands. The child *device-classes* extend the top level *device-class*, respond to all genetic commands as well as their instrument-specific commands. New commands can be easily defined and implemented for a specific experimental application by extending *device-class* child. For all commands above, use MQTT topic: UUID/device name.

## Footnotes

1 https://github.com/braingeneers/integrated-system-v1-paper

2 https://github.com/braingeneers/braingeneerspy

## References

[1] Kelley, K.W., Pas, ca, S.P.: Human brain organogenesis: Toward a cellular understanding of development and disease. Cell 185(1), 42–61 (2022) 10.1016/j.cell.2021.10.003. Accessed 2024-02-13

[2] Eiraku, M., Watanabe, K., Matsuo-Takasaki, M., Kawada, M., Yonemura, S., Matsumura, M., Wataya, T., Nishiyama, A., Muguruma, K., Sasai, Y.: Self-Organized Formation of Polarized Cortical Tissues from ESCs and Its Active Manipulation by Extrinsic Signals. Cell Stem Cell 3(5), 519–532 (2008) 10.1016/j.stem.2008.09.002. Publisher: Elsevier. Accessed 2020-04-14

[3] Lancaster, M.A., Renner, M., Martin, C.-A., Wenzel, D., Bicknell, L.S., Hurles, M.E., Homfray, T., Penninger, J.M., Jackson, A.P., Knoblich, J.A.: Cerebral organoids model human brain development and microcephaly. Nature 501(7467), 373–379 (2013) 10.1038/nature12517. Number: 7467 Publisher: Nature Publishing Group. Accessed 2021-02-04

[4] Pollen, A.A., Bhaduri, A., Andrews, M.G., Nowakowski, T.J., Meyerson, O.S., Mostajo-Radji, M.A., Lullo, E.D., Alvarado, B., Bedolli, M., Dougherty, M.L., Fiddes, I.T., Kronenberg, Z.N., Shuga, J., Leyrat, A.A., West, J.A., Bershteyn, M., Lowe, C.B., Pavlovic, B.J., Salama, S.R., Haussler, D., Eichler, E.E., Kriegstein, A.R.: Establishing Cerebral Organoids as Models of Human-Specific Brain Evolution. Cell 176(4), 743–75617 (2019) 10.1016/j.cell.2019.01.017. Publisher: Elsevier. Accessed 2021-02-03

[5] Giandomenico, S.L., Mierau, S.B., Gibbons, G.M., Wenger, L.M.D., Masullo, L., Sit, T., Sutcliffe, M., Boulanger, J., Tripodi, M., Derivery, E., Paulsen, O., Lakatos, A., Lancaster, M.A.: Cerebral organoids at the air–liquid interface generate diverse nerve tracts with functional output. Nature Neuroscience 22(4), 669–679 (2019) 10.1038/s41593-019-0350-2. Accessed 2019-09-29

[6] Andersen, J., Revah, O., Miura, Y., Thom, N., Amin, N.D., Kelley, K.W., Singh, M., Chen, X., Thete, M.V., Walczak, E.M., Vogel, H., Fan, H.C., Pasca, S.P.: Generation of Functional Human 3D Cortico-Motor Assembloids. Cell 183(7), 1913–192926 (2020) 10.1016/j.cell.2020.11.017. Accessed 2021-04-13

[7] Gonzalez, C., Armijo, E., Bravo-Alegria, J., Becerra-Calixto, A., Mays, C.E., Soto, C.: Modeling amyloid beta and tau pathology in human cerebral organoids. Molecular Psychiatry 23(12), 2363–2374 (2018) 10.1038/s41380-018-0229-8. Publisher: Nature Publishing Group. Accessed 2024-03-07

[8] Li, C., Fleck, J.S., Martins-Costa, C., Burkard, T.R., Themann, J., Stuempflen, M., Peer, A.M., Vertesy Littleboy, J.B., Esk, C., Elling, U., Kasprian, G., Corsini, N.S., Treutlein, B., Knoblich, J.A.: Single-cell brain organoid screening identifies developmental defects in autism. Nature 621(7978), 373–380 (2023) 10.1038/s41586-023-06473-y. Number: 7978 Publisher: Nature Publishing Group. Accessed 2024-02-13

[9] Miura, Y., Li, M.-Y., Revah, O., Yoon, S.-J., Narazaki, G., Pasca, S.P.: Engineering brain assembloids to interrogate human neural circuits. Nature Protocols 17(1), 15–35 (2022) 10.1038/s41596-021-00632-z. Number: 1 Publisher: Nature Publishing Group. Accessed 2024-02-13

[10] Sloan, S.A., Darmanis, S., Huber, N., Khan, T.A., Birey, F., Caneda, C., Reimer, R., Quake, S.R., Barres, B.A., Pasca, S.P.: Human Astrocyte Maturation Captured in 3D Cerebral Cortical Spheroids Derived from Pluripotent Stem Cells. Neuron 95(4), 779–7906 (2017) 10.1016/j.neuron.2017.07.035. Accessed 2024-02-13

[11] Passaro, A.P., Stice, S.L.: Electrophysiological Analysis of Brain Organoids: Current Approaches and Advancements. Frontiers in Neuroscience 14 (2021) 10.3389/fnins.2020.622137. Publisher: Frontiers. Accessed 2021-06-11

[12] Voitiuk, K., Geng, J., Keefe, M.G., Parks, D.F., Sanso, S.E., Hawthorne, N., Freeman, D.B., Mostajo-Radji, M.A., Nowakowski, T.J., Salama, S.R., Teodorescu, M., Haussler, D.: Light-weight Electrophysiology Hardware and Software Platform for Cloud-Based Neural Recording Experiments. bioRxiv, 2021–0518444685 (2021) 10.1101/2021.05.18.444685. Publisher: Cold Spring Harbor Laboratory Section: New Results. Accessed 2021-05-21

[13] Bertoncello, I.: Optimizing the Cell Culture Microenvironment. In: Bertoncello, I. (ed.) Mouse Cell Culture: Methods and Protocols. Methods in Molecular Biology, pp. 23–30. Springer, New York, NY (2019). 10.1007/978-1-4939-9086-32 Accessed 2024-02-13

[14] Walters, E.A., Brown, J.L., Krisher, R., Voelkel, S., Swain, J.E.: Impact of a controlled culture temperature gradient on mouse embryo development and morphokinetics. Reproductive BioMedicine Online 40(4), 494–499 (2020) 10.1016/j.rbmo.2019.12.015. Accessed 2024-02-13

[15] Kong, F., Yuan, L., Zheng, Y.F., Chen, W.: Automatic liquid handling for life science: a critical review of the current state of the art. Journal of Laboratory Automation 17(3), 169–185 (2012) 10.1177/2211068211435302

[16] Torres-Acosta, M.A., Lye, G.J., Dikicioglu, D.: Automated liquid-handling operations for robust, resilient, and efficient bio-based laboratory practices. Biochemical Engineering Journal 188, 108713 (2022) 10.1016/j.bej.2022.108713. Accessed 2024-03-13

[17] Waheed, S., Cabot, J.M., Macdonald, N.P., Lewis, T., Guijt, R.M., Paull, B., Breadmore, M.C.: 3D printed microfluidic devices: enablers and barriers. Lab on a Chip 16(11), 1993–2013 (2016) 10.1039/C6LC00284F. Publisher: The Royal Society of Chemistry. Accessed 2024-03-13

[18] Macdonald, N.P., Cabot, J.M., Smejkal, P., Guijt, R.M., Paull, B., Breadmore, M.C.: Comparing Microfluidic Performance of Three-Dimensional (3D) Printing Platforms. Analytical Chemistry 89(7), 3858–3866 (2017) 10.1021/acs.analchem.7b00136. Publisher: American Chemical Society. Accessed 2024-03-13

[19] Hu, D., Gong, Y., Seibel, E.J., Sekhar, L.N., Hannaford, B.: Semi-autonomous image-guided brain tumour resection using an integrated robotic system: A bench-top study. The International Journal of Medical Robotics and Computer Assisted Surgery 14(1), 1872 (2018) 10.1002/rcs.1872. eprint: https://onlinelibrary.wiley.com/doi/pdf/10.1002/rcs.1872. Accessed 2024-03-13

[20] Slattery, A., Wen, Z., Tenblad, P., Sanjosé-Orduna, J., Pintossi, D., Hartog, T., Noël, T.: Automated self-optimization, intensification, and scale-up of photocatalysis in flow. Science 383(6681), 1817 (2024) 10.1126/science.adj1817. Publisher: American Association for the Advancement of Science. Accessed 2024-03-07

[21] Konishi, S., Hashimoto, T., Nakabuchi, T., Ozeki, T., Kajita, H.: Cell and tissue system capable of automated culture, stimulation, and monitor with the aim of feedback control of organs-on-a-chip. Scientific Reports 11(1), 2999 (2021) 10.1038/s41598-020-80447-2. Publisher: Nature Publishing Group. Accessed 2024-03-07

[22] Chen, Z., Blair, G.J., Cao, C., Zhou, J., Aharoni, D., Golshani, P., Blair, H.T., Cong, J.: FPGA-Based In-Vivo Calcium Image Decoding for Closed-Loop Feedback Applications. IEEE Transactions on Biomedical Circuits and Systems 17(2), 169–179 (2023) 10.1109/TBCAS.2023.3268130. Conference Name: IEEE Transactions on Biomedical Circuits and Systems. Accessed 2024-03-14

[23] Passos, J., Lopes, S.I., Clemente, F.M., Moreira, P.M., Rico-González, M., Bezerra, P., Rodrigues, L.P.: Wearables and Internet of Things (IoT) Technologies for Fitness Assessment: A Systematic Review. Sensors 21(16), 5418 (2021) 10.3390/s21165418. Number: 16 Publisher: Multidisciplinary Digital Publishing Institute. Accessed 2024-02-13

[24] Cariou, C., Moiroux-Arvis, L., Pinet, F., Chanet, J.-P.: Internet of Underground Things in Agriculture 4.0: Challenges, Applications and Perspectives. Sensors 23(8), 4058 (2023) 10.3390/s23084058. Number: 8 Publisher: Multidisciplinary Digital Publishing Institute. Accessed 2024-02-13

[25] Su, P., Chen, Y., Lu, M.: Smart city information processing under internet of things and cloud computing. The Journal of Supercomputing 78(3), 3676–3695 (2022) 10.1007/s11227-021-03972-5. Accessed 2024-02-13

[26] Sadhu, P.K., Yanambaka, V.P., Abdelgawad, A.: Internet of Things: Security and Solutions Survey. Sensors 22(19), 7433 (2022) 10.3390/s22197433. Number: 19 Publisher: Multidisciplinary Digital Publishing Institute. Accessed 2024-02-13

[27] Kelly, J.T., Campbell, K.L., Gong, E., Scuffham, P.: The Internet of Things: Impact and Implications for Health Care Delivery. Journal of Medical Internet Research 22(11), 20135 (2020) 10.2196/20135. Company: Journal of Medical Internet Research Distributor: Journal of Medical Internet Research Institution: Journal of Medical Internet Research Label: Journal of Medical Internet Research Publisher: JMIR Publications Inc., Toronto, Canada. Accessed 2024-02-13

[28] Parks, D.F., Voitiuk, K., Geng, J., Elliott, M.A.T., Keefe, M.G., Jung, E.A., Robbins, A., Baudin, P.V., Ly, V.T., Hawthorne, N., Yong, D., Sanso, S.E., Rezaee, N., Sevetson, J.L., Seiler, S.T., Currie, R., Pollen, A.A., Hengen, K.B., Nowakowski, T.J., Mostajo-Radji, M.A., Salama, S.R., Teodorescu, M., Haussler, D.: IoT cloud laboratory: Internet of Things architecture for cellular biology. Internet of Things, 100618 (2022) 10.1016/j.iot.2022.100618. Accessed 2022-09-28

[29] Müller, J., Ballini, M., Livi, P., Chen, Y., Radivojevic, M., Shadmani, A., Viswam, V., Jones, I.L., Fiscella, M., Diggelmann, R., Stettler, A., Frey, U., Bakkum, D.J., Hierlemann, A.: High-resolution CMOS MEA platform to study neurons at subcellular, cellular, and network levels. Lab on a chip 15(13), 2767–2780 (2015) 10.1039/c5lc00133a. Accessed 2024-02-13

[30] Seiler, S.T., Mantalas, G.L., Selberg, J., Cordero, S., Torres-Montoya, S., Baudin, P.V., Ly, V.T., Amend, F., Tran, L., Hoffman, R.N., Rolandi, M., Green, R.E., Haussler, D., Salama, S.R., Teodorescu, M.: Modular automated microfluidic cell culture platform reduces glycolytic stress in cerebral cortex organoids. bioRxiv. Pages: 2022.07.13.499938 Section: New Results (2022). 10.1101/2022.07.13.499938. https://www.biorxiv.org/content/10.1101/2022.07. 13.499938v1 Accessed 2022-07-24

[31] Denninger, J.K., Chen, X., Turkoglu, A.M., Sarchet, P., Volk, A.R., Rieskamp, J.D., Yan, P., Kirby, E.D.: Defining the adult hippocampal neural stem cell secretome: In vivo versus in vitro transcriptomic differences and their correlation to secreted protein levels. Brain Research 1735, 146717 (2020) 10.1016/j.brainres.2020.146717. Accessed 2024-02-13

[32] Watanabe, M., Buth, J.E., Vishlaghi, N., Torre-Ubieta, L., Taxidis, J., Khakh, B.S., Coppola, G., Pearson, C.A., Yamauchi, K., Gong, D., Dai, X., Damoiseaux, R., Aliyari, R., Liebscher, S., Schenke-Layland, K., Caneda, C., Huang, E.J., Zhang, Y., Cheng, G., Geschwind, D.H., Golshani, P., Sun, R., Novitch, B.G.: Self-Organized Cerebral Organoids with Human-Specific Features Predict Effective Drugs to Combat Zika Virus Infection. Cell Reports 21(2), 517–532 (2017) 10.1016/j.celrep.2017.09.047. Accessed 2024-02-13

[33] Ojala, M., Garriga, G.C.: Permutation Tests for Studying Classifier Performance. The Journal of Machine Learning Research 11, 1833–1863 (2010)

[34] Elliott, M.A.T., Schweiger, H.E., Robbins, A., Vera-Choqqueccota, S., Ehrlich, D., Hernandez, S., Voitiuk, K., Geng, J., Sevetson, J.L., Core, C., Rosen, Y.M., Teodorescu, M., Wagner, N.O., Haussler, D., Mostajo-Radji, M.A.: Internet-Connected Cortical Organoids for Project-Based Stem Cell and Neuroscience Education. eNeuro 10(12) (2023) 10.1523/ENEURO.0308-23. 2023. Publisher: Society for Neuroscience Section: Research Article: New Research. Accessed 2024-01-11

[35] Park, Y., Hernandez, S., Hernandez, C.O., Schweiger, H.E., Li, H., Voitiuk, K., Dechiraju, H., Hawthorne, N., Muzzy, E.M., Selberg, J.A., Sullivan, F.N., Urcuyo, R., Salama, S.R., Aslankoohi, E., Knight, H.J., Teodorescu, M., Mostajo-Radji, M.A., Rolandi, M.: Modulation of neuronal activity in cortical organoids with bioelectronic delivery of ions and neurotransmitters. Cell Reports Methods 4(1), 100686 (2024) 10.1016/j.crmeth.2023.100686. Accessed 2024-01-30

[36] Chen, Y., Austin, R.H., Sturm, J.C.: On-chip cell labelling and washing by capture and release using microfluidic trap arrays. Biomicrofluidics 11(5), 054107 (2017) 10.1063/1.4985771. Accessed 2024-02-13

[37] Lu, X., Ai, Y.: Automatic Microfluidic Cell Wash Platform for Purifying Cells in Suspension: Puriogen. Analytical Chemistry 94(26), 9424–9433 (2022) 10.1021/acs.analchem.2c01616. Publisher: American Chemical Society. Accessed 2024-02-13

[38] Lee, C.-S.: Grand Challenges in Microfluidics: A Call for Biological and Engineering Action. Frontiers in Sensors 1 (2020). Accessed 2024-02-13

[39] Pachitariu, M., Steinmetz, N., Kadir, S., Carandini, M. D H.K.: Kilosort: realtime spike-sorting for extracellular electrophysiology with hundreds of channels. bioRxiv, 061481 (2016) 10.1101/061481. Publisher: Cold Spring Harbor Laboratory Section: New Results. Accessed 2020-03-17

[40] Fair, S.R., Julian, D., Hartlaub, A.M., Pusuluri, S.T., Malik, G., Summerfied, T.L., Zhao, G., Hester, A.B., Ackerman, W.E., Hollingsworth, E.W., Ali, M., McElroy, C.A., Buhimschi, I.A., Imitola, J., Maitre, N.L., Bedrosian, T.A., Hester, M.E.: Electrophysiological Maturation of Cerebral Organoids Correlates with Dynamic Morphological and Cellular Development. Stem Cell Reports 15(4), 855–868 (2020) 10.1016/j.stemcr.2020.08.017. Accessed 2021-01-25

[41] Schröter, M., Wang, C., Terrigno, M., Hornauer, P., Huang, Z., Jagasia, R., Hierlemann, A.: Functional imaging of brain organoids using high-density microelectrode arrays. Mrs Bulletin 47(6), 530–544 (2022) 10.1557/s43577-022-00282-w. Accessed 2024-02-13

[42] Sharf, T., Molen, T., Glasauer, S.M.K., Guzman, E., Buccino, A.P., Luna, G., Cheng, Z., Audouard, M., Ranasinghe, K.G., Kudo, K., Nagarajan, S.S., Tovar, K.R., Petzold, L.R., Hierlemann, A., Hansma, P.K., Kosik, K.S.: Functional neuronal circuitry and oscillatory dynamics in human brain organoids. Nature Communications 13(1), 4403 (2022) 10.1038/s41467-022-32115-4. Number: 1 Publisher: Nature Publishing Group. Accessed 2022-07-29

[43] Chini, M., Hnida, M., Kostka, J.K., Chen, Y.-N., Hanganu-Opatz, I.L.: Preconfigured architecture of the developing mouse brain. Cell Reports 43(6) (2024) 10.1016/j.celrep.2024.114267. Publisher: Elsevier. Accessed 2024-12-01

[44] Rzechorzek, N.M., Sutcliffe, M.A., Mihut, A., Baranes, K., Karam, N., Sánchez, D.L.-D., Peak-Chew, S.Y., Zeng, A., Poulin, N., Seinkmane, E., Karim, K., Proctor, C.M., Kotter, M., Lancaster, M.A., Beale, A.D.: Circadian clocks in human cerebral organoids. bioRxiv. Pages: 2024.02.20.580978 Section: New Results (2024). 10.1101/2024.02.20.580978. https://www.biorxiv.org/content/10.1101/2024.02.20.580978v1 Accessed 2024-03-04

[45] Lee, S., Hong, C.I.: Organoids as Model Systems to Investigate Circadian Clock-Related Diseases and Treatments. Frontiers in Genetics 13, 874288 (2022) 10.3389/fgene.2022.874288. Accessed 2024-03-14

[46] Turrigiano, G.G., Nelson, S.B.: Homeostatic plasticity in the developing nervous system. Nature Reviews Neuroscience 5(2), 97–107 (2004) 10.1038/nrn1327. Number: 2 Publisher: Nature Publishing Group. Accessed 2024-02-13

[47] Urenda, J.-P., Del Dosso, A., Birtele, M., Quadrato, G.: Present and Future Modeling of Human Psychiatric Connectopathies With Brain Organoids. Biological Psychiatry 93(7), 606–615 (2023) 10.1016/j.biopsych.2022.12.017. Accessed 2024-02-13

[48] Kirillov, A., Mintun, E., Ravi, N., Mao, H., Rolland, C., Gustafson, L., Xiao, T., Whitehead, S., Berg, A.C., Lo, W.-Y., Dollár, P., Girshick, R.: Segment Anything. arXiv. arXiv:2304.02643 [cs] (2023). 10.48550/arXiv.2304.02643. http://arxiv.org/abs/2304.02643 Accessed 2024-03-14

[49] Ballini, M., Müller, J., Livi, P., Chen, Y., Frey, U., Stettler, A., Shadmani, A., Viswam, V., Jones, I.L., Jäckel, D., Radivojevic, M., Lewandowska, M.K., Gong, W., Fiscella, M., Bakkum, D.J., Heer, F., Hierlemann, A.: A 1024-Channel CMOS Microelectrode Array With 26,400 Electrodes for Recording and Stimulation of Electrogenic Cells In Vitro. IEEE Journal of Solid-State Circuits 49(11), 2705–2719 (2014) 10.1109/JSSC.2014.2359219. Conference Name: IEEE Journal of Solid-State Circuits. Accessed 2024-02-13

[50] National Research Platform. https://nationalresearchplatform.org/ Accessed 2024-02-13

[51] Park, Y., Hernandez, S., Hernandez, C.O., Schweiger, H.E., Li, H., Voitiuk, K., Dechiraju, H., Hawthorne, N., Muzzy, E.M., Selberg, J.A., Sullivan, F.N., Urcuyo, R., Salama, S.R., Aslankoohi, E., Teodorescu, M., Mostajo-Radji, M.A., Rolandi, M.: Modulation of neuronal activity in cortical organoids with bioelectronic delivery of ions and neurotransmitters. bioRxiv. Pages: 2023.06.10.544416 Section: New Results (2023). 10.1101/2023.06.10.544416. https://www.biorxiv.org/content/10.1101/2023.06.10.544416v1 Accessed 2023-07-05

[52] Hill, D.N., Mehta, S.B., Kleinfeld, D.: Quality Metrics to Accompany Spike Sorting of Extracellular Signals. Journal of Neuroscience 31(24), 8699–8705 (2011) 10.1523/JNEUROSCI.0971-11.2011. Publisher: Society for Neuroscience Section: Toolbox. Accessed 2023-05-15

[53] Slack: Unlock your productivity potential with Slack Platform. https://slack.com Accessed 2024-03-14

[54] Potter, S.M., DeMarse, T.B.: A new approach to neural cell culture for long-term studies. Journal of Neuroscience Methods 110(1), 17–24 (2001) 10.1016/S0165-0270(01)00412-5. Accessed 2024-02-22

