## Supplemental Methods for "A feedback-driven brain organoid platform enables automated maintenance and high-resolution neural activity monitoring"

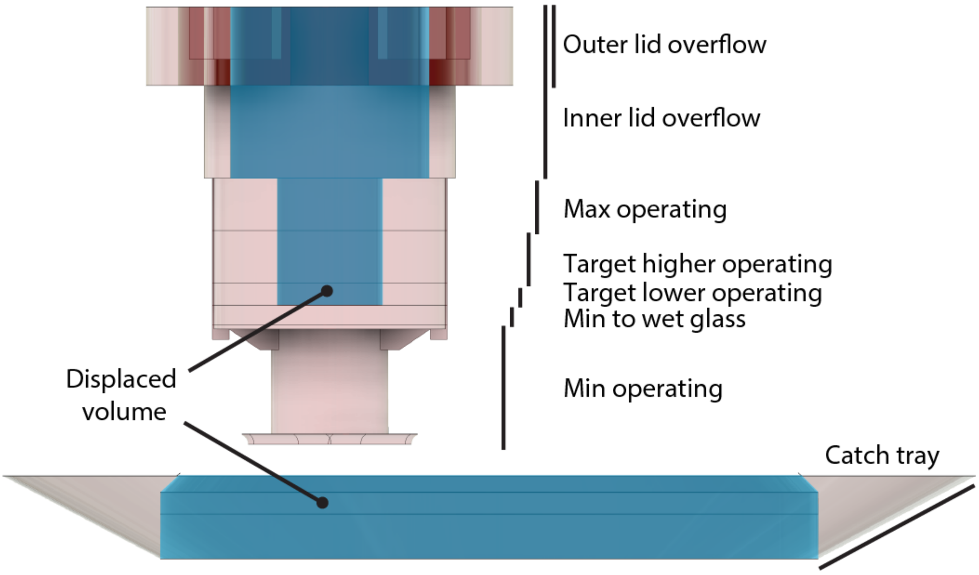


**Supplementary Fig. 1  Diagram of operating ranges of the microfluidic culture chamber.** Shaded pink areas represent volumes where media is collected. Shaded blue areas mark displaced volumes (where there is no media stored). The numerical volumes for each operating range are listed in Supplementary Table 1.

| Feature | Delta (µL) | Running Total (µL) |
|---|---|---|
| Min operating | 172.2 | 172.2 |
| Min to wet glass | 94.0 | 266.2 |
| Target lower operating | 88.1 | 354.3 |
| Target higher operating | 204.9 | 559.2 |
| Max operating | 204.9 | 764.1 |
| **Total operating capacity:** | | **764.1** |
| Inner lid overflow | 345.1 | 1109.2 |
| Outer lid overflow | 464.1 | 1573.3 |
| **Total chip capacity:** | | **1573.3** |
| Catch tray | 1539.5 | 3112.8 |
| **Total overflowed capacity:** | | **3112.8** |

**Supplementary Table. 1** Numerical operating volume ranges based on the microfluidic culture chamber's 3D model (CAD) measurements. Illustrations of operating ranges are shown in Supplementary Figure 1. The Feature column lists critical points in the microfluidic culture chamber. The Delta column is the volume space between each feature, and the Running Total column is the volume from the floor to the feature.

| State | Description |
|---|---|
| **SHUTDOWN** | The device has been turned off gracefully and won't respond until it's turned back on. |
| **IDLE** | The device is not assigned to any experiments and not doing anything at the moment, and is missing physical prerequisites (i.e., a reagent or piece of hardware) to be able to perform its job. |
| **PRIMED** | The device is not assigned to any experiments and not doing anything at the moment, but it has all the physical prerequisites to perform its job. |
| **READY** | The device is assigned to an experiment and is ready to execute a command. |
| **WAITING** | The device has received a command to PAUSE and is waiting until a given time to resume performing jobs. |
| **EXEC** | The device is actively executing a job command. |

**Supplementary Table. 1 Device states.** The *device-class* is structured as a finite-state machine, with a defined set of states (SHUTDOWN, IDLE, PRIMED, READY, PAUSED, EXEC) that describe its status. The finite-state machine reads a set of inputs and changes to a different state based on those inputs. The inputs can be user physical interactions (i.e., button press, linkage of consumables, etc.), MQTT messages containing job requests, or scheduled events.

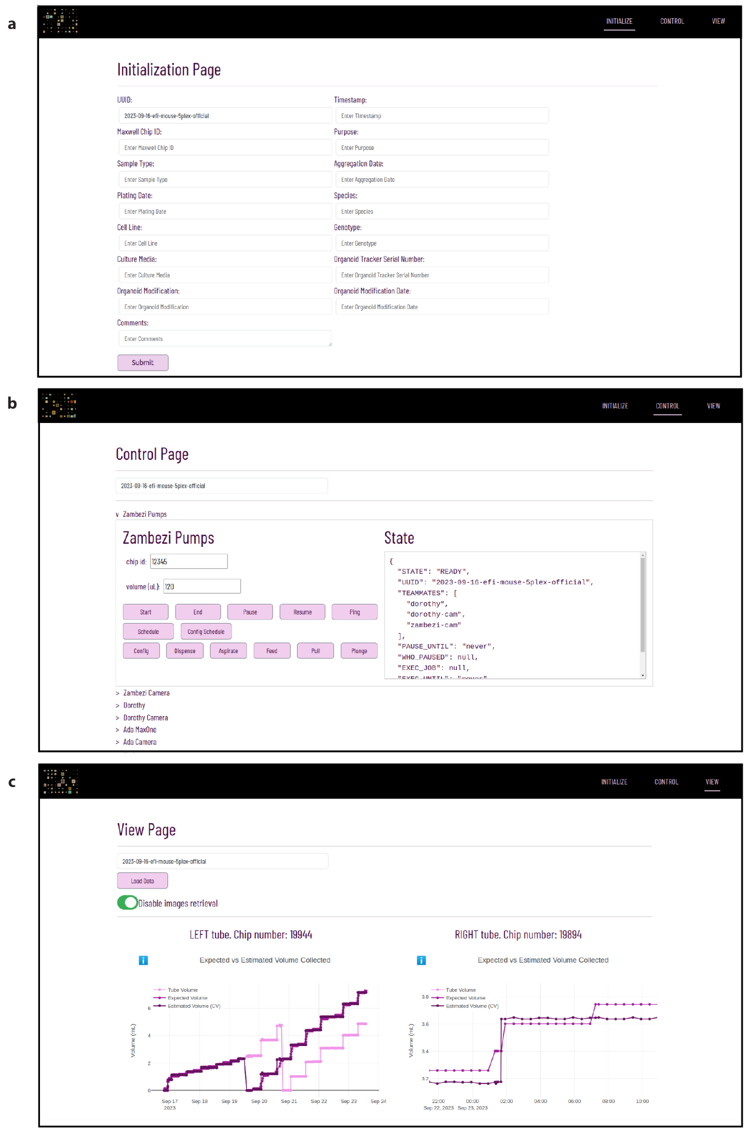


**Supplementary Fig. 2 Webpage user interface screenshots. (a)** Initialization page: Users can input details about the experiment and the biological samples. **(b)** Control page: Users can access and control every device involved in the experiment. **(c)** Visualization page: It includes three graph types. (1) Expected versus Estimated Volume Graph: compare volumes determined by the computer vision algorithm with volume according to pump metrics, highlighting any discrepancies and mismatching data. (2) Expected minus Estimated Graph: It shows the difference between the pump metrics and computer vision estimates for each device. They are designed to quickly identify alignment or discrepancies between these two methods, where values close to zero suggest good alignment, and deviations indicate measurement inaccuracies. (3) Collected Volume According to Computer Vision and Pump Graph: This graph contrasts the volume of media collected as reported by the pump system with that detected by the Computer Vision algorithm, which is crucial for assessing feeding accuracy. For example, if the pump indicates a feed of 300 µL, but the Computer Vision only detects 150 µL, this discrepancy is highlighted.

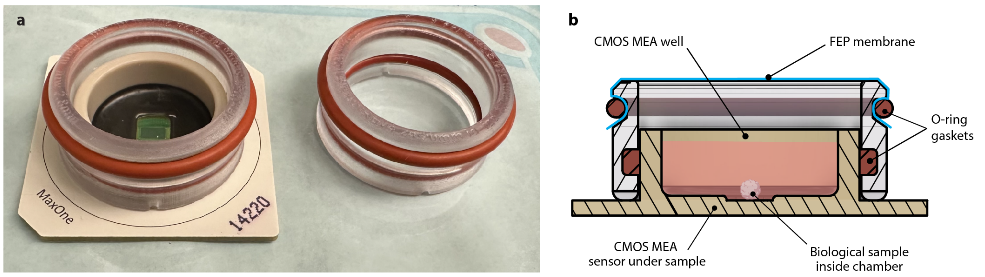


**Supplementary Fig. 3  3D printed breathable membrane lid used for Controls modeled after designs by Potter [? ].** **(a)** Picture of the membrane lid and HD-MEA. The chamber is comprised of biocompatible 3D-printed parts, sealed by O-rings to the HD-MEA, and imaged through the FEP membrane stretched over the top with an O-ring. **(b)** Cross-sectional rendering depicting the fluid path and position of the sample.

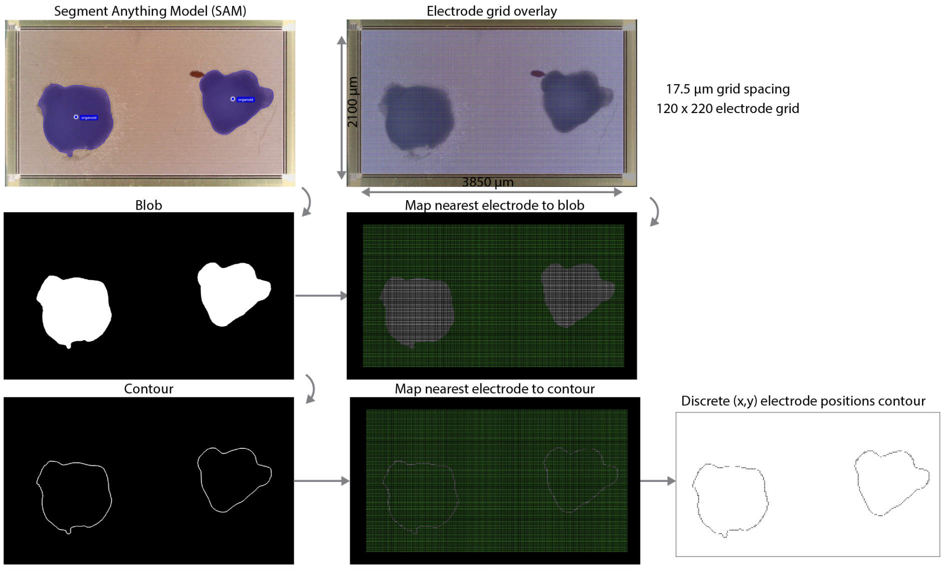


**Supplementary Fig. 4**  Organoid boundary segmentation process.

.

| Command | Description |
|---|---|
| START* | Initiates an experiment process on a specified device. The device must not be engaged in another experiment and should be in a PRIMED state. If successful, the device acknowledges the request and changes its state to READY. |
| END† | Ends an ongoing experiment on a device or all devices associated with an experiment UUID. The device(s) will drop current tasks and reset to the IDLE state. |
| STATUS† | Retrieves the current status, state, associated experiment UUID, teammates, and job schedule. Works in any device state. |
| PAUSE† | Temporarily halts the device's ability to start working on new commands for a specified duration. If the device is already paused or not part of an experiment, it will return an error. Otherwise, the device will successfully change its state to WAITING. |
| RESUME† | Requests the device to continue execution after a pause. Only the device that initiated the pause can send a resume command. The PAUSED device will successfully change its state to WAITING. |
| SCHEDULE† | Adds, clears, or retrieves scheduled tasks for the device. The device will execute the specified task payload at the specified time every X hours or minutes (unless it's WAITING, then it will do backlogged tasks at the easiest convenience). Works in any state. |
| STOP† | Requests the cancellation of a running task. If there is no task running, an error message is returned. |
| PING† | Requests the device to respond with a ping message. This is used to check if the device is online and listening to a given topic. Works in any state. |
| SLACK‡ | Posts a message to Slack. The message can contain text and/or an image. |

**Supplementary Table. 1 Generic commands.** The parent *device-class* responds to a generic set of commands. Commands are sent on hierarchical MQTT topics that allow widening and narrowing of scope. We used each experiment's Universal Unique Identifier (UUID) and each device's name as part of the topic. If a device is not part of an experiment, the default UUID is NONE.
* Use MQTT topic: `NONE/device` because no experiment assigned yet
† Use MQTT topic: `UUID/device` or just `UUID` to address all
‡ Use MQTT topic: `TOSLACK`

| Command | Device | Description |
|---|---|---|
| RECORD | HD-MEA | Performs an electrophysiology recording for a defined period of time. |
| PICTURE | Camera(s) | Takes a picture from the camera. |
| FEED | Pump | Performs a cycle of aspirating spent and dispensing fresh media of the pre-configured volume. |
| ASPIRATE | Pump | Aspirates a specified volume of liquid (mL) from the culture chamber. |
| DISPENSE | Pump | Dispenses a specified volume of liquid (mL) to the culture chamber. |
| PULL | Pump | A rapid, full-syringe aspiration to assist pulling media through high resistance or clogs. |
| SPIKESORT | Spike sorting | Spike sorts a specified dataset stored in S3 using the analysis pipeline. |
| ESTIMATE | Estimator | Estimates the amount (mL) of media in a collection reservoir by applying computer vision analysis to a specified image of the collection reservoir stored in S3. |

**Supplementary Table. 1 Application-specific commands.** The child *device-classes* extend the top level *device-class*, respond to all genetic commands as well as their instrument-specific commands. New commands can be easily defined and implemented for a specific experimental application by extending *device-class* child. For all commands above, use MQTT topic: `UUID/device_name`.
